# Genus-wide characterization of bumblebee genomes reveals variation associated with key ecological and behavioral traits of pollinators

**DOI:** 10.1101/2020.05.29.122879

**Authors:** Cheng Sun, Jiaxing Huang, Yun Wang, Xiaomeng Zhao, Long Su, Gregg W.C. Thomas, Mengya Zhao, Xingtan Zhang, Irwin Jungreis, Manolis Kellis, Saverio Vicario, Igor V. Sharakhov, Semen M. Bondarenko, Martin Hasselmann, Chang N Kim, Benedict Paten, Luca Penso-Dolfin, Li Wang, Yuxiao Chang, Qiang Gao, Ling Ma, Lina Ma, Zhang Zhang, Hongbo Zhang, Huahao Zhang, Livio Ruzzante, Hugh M. Robertson, Yihui Zhu, Yanjie Liu, Huipeng Yang, Lele Ding, Quangui Wang, Weilin Xu, Cheng Liang, Michael W. Itgen, Lauren Mee, Ben M. Sadd, Gang Cao, Ze Zhang, Matthew Hahn, Sarah Schaack, Seth M. Barribeau, Paul H. Williams, Robert M. Waterhouse, Rachel Lockridge Mueller

## Abstract

Bumblebees are a diverse group of globally important pollinators in natural ecosystems and for agricultural food production. With both eusocial and solitary lifecycle phases, and some social parasite species, they are especially interesting models to understand social evolution, behavior, and ecology. Reports of many species in decline point to pathogen transmission, habitat loss, pesticide usage, and global climate change, as interconnected causes. These threats to bumblebee diversity make our reliance on a handful of well-studied species for agricultural pollination particularly precarious. To broadly sample bumblebee genomic and phenotypic diversity, we *de novo* sequenced and assembled the genomes of 17 species, representing all 15 subgenera, producing the first genus-wide quantification of genetic and genomic variation potentially underlying key ecological and behavioral traits. The species phylogeny resolves subgenera relationships while incomplete lineage sorting likely drives high levels of gene tree discordance. Five chromosome-level assemblies show a stable 18-chromosome karyotype, with major rearrangements creating 25 chromosomes in social parasites. Differential transposable element activity drives changes in genome sizes, with putative domestications of repetitive sequences influencing gene coding and regulatory potential. Dynamically evolving gene families and signatures of positive selection point to genus-wide variation in processes linked to foraging, diet and metabolism, immunity and detoxification, as well as adaptations for life at high altitudes. These high-quality genomic resources capture natural genetic and phenotypic variation across bumblebees, offering new opportunities to advance our understanding of their remarkable ecological success and to identify and manage current and future threats.

## Introduction

Bumblebees (Hymenoptera: *Apidae)* are a group of pollinating insects comprising the genus *Bombus,* which are economically important for crop pollination (Garibaldi, et al. 2013; Martin, et al. 2019; Velthuis and Van Doorn 2006). Bumblebees are also ecologically important pollinators, serving as the sole or predominant pollinators of many wild plants (Fontaine, et al. 2005; Goulson, et al. 2008). They are particularly charismatic social insects that exhibit complex behaviors such as learning through observation (Alem, et al. 2016) and damaging leaves to stimulate earlier flowering (Pashalidou, et al. 2020). Global and local environmental changes have resulted in some species declining in abundance and others remaining stable or even increasing (Bartomeus, et al. 2013; Cameron, et al. 2011; Cameron and Sadd 2019; Koch, et al. 2015). Decline in bumblebee abundance and distribution resulting from habitat loss, pathogen transmission, climate change, and agrochemical exposure is threatening pollination services to both wild plants and crops, raising concerns for bumblebees, the plant species they service, food security, and ecosystem stability (Cameron and Sadd 2019; Goulson, et al. 2015; Grixti, et al. 2009; Potts, et al. 2010; Soroye, et al. 2020; Williams and Osborne 2009).

Bumblebees comprise ~250 extant species classified into 15 subgenera (Williams 1998; Williams, et al. 2018). The initial diversification of *Bombus* lineages occurred ~25-40 million years ago (Ma), near the Eocene-Oligocene boundary ~34 Ma (Hines 2008; Williams 1998). Bumblebees display considerable interspecific diversity in morphology, food preference, pathogen incidence, and exhibit diverse life histories and ecologies (Arbetman, et al. 2017; Persson, et al. 2015; Sikora and Kelm 2012; Williams 1994). Members of the subgenus *Mendacibombus,* the sister group to all other extant bumblebees, are high-elevation specialists with distributions centered on the Qinghai-Tibetan plateau (Williams, et al. 2018). Species in the subgenus *Psithyrus* exhibit social parasitism; they do not have a worker caste, and they feed on food collected by host workers (Lhomme and Hines 2019). Bumblebees are distributed across the globe, from Greenland to the Amazon Basin and from sea level to altitudes of 5,640 m in the Himalayas, where they occupy diverse habitats, from alpine meadows to lowland tropical forest (Williams and Paul 1985; Williams, et al. 2018). Much remains to be learned about bumblebees. For example, little is known about the underlying genetic and genomic variation that gives rise to these diverse phenotypes, including their differential responses to changing environments.

To broadly sample this genomic and phenotypic diversity, we performed *de novo* sequencing and assembly of the genomes of 17 bumblebee species, representing all of the 15 subgenera within the genus *Bombus*. Integrating these datasets with two previously published bumblebee genomes, we performed comparative analyses of genome structures, genome contents, and gene evolutionary dynamics across the phylogeny. Our results characterizing bumblebee gene and genome evolution provide the first genus-wide quantification of genetic and genomic variation potentially underlying key eco-ethological traits.

## Results

### High quality genomic resources for all 15 *Bombus* subgenera

Sequencing and assembly strategies resulted in high quality genomic resources with 12 scaffold-level and five chromosome-level genome assemblies (Table 1). Criteria including phylogenetic position, species traits, and geographic distribution were applied to select species for whole genome sequencing from across the genus. For the five species for which sufficient samples could be collected, high-throughput chromatin conformation capture (Hi-C) (Belton, et al. 2012) was used to produce chromosome-level genome assemblies (Table 1). A total of 17 species were selected (Additional file 1: Table S1), which span all 15 subgenera within *Bombus* (Williams, et al. 2008). Among these, two species *(B. superbus* and *B. waltoni)* are from *Mendacibombus,* the earliest split in the *Bombus* phylogeny; four species (*B. superbus, B. waltoni, B. skorikovi,* and *B. difficillimus)* inhabit high elevations (> 4000 m above sea level); two species (*B. turneri* and *B. skorikovĩ)* exhibit social parasitism; and one species (*B. polaris*) is endemic to Arctic/subarctic regions (Williams, et al. 2019). In addition, species traits including range size, tongue length, parasite incidence, and decline status vary across the selected species (Arbetman, et al. 2017; Williams 1994)(Additional file 1: Table S1).

**Table 1.**
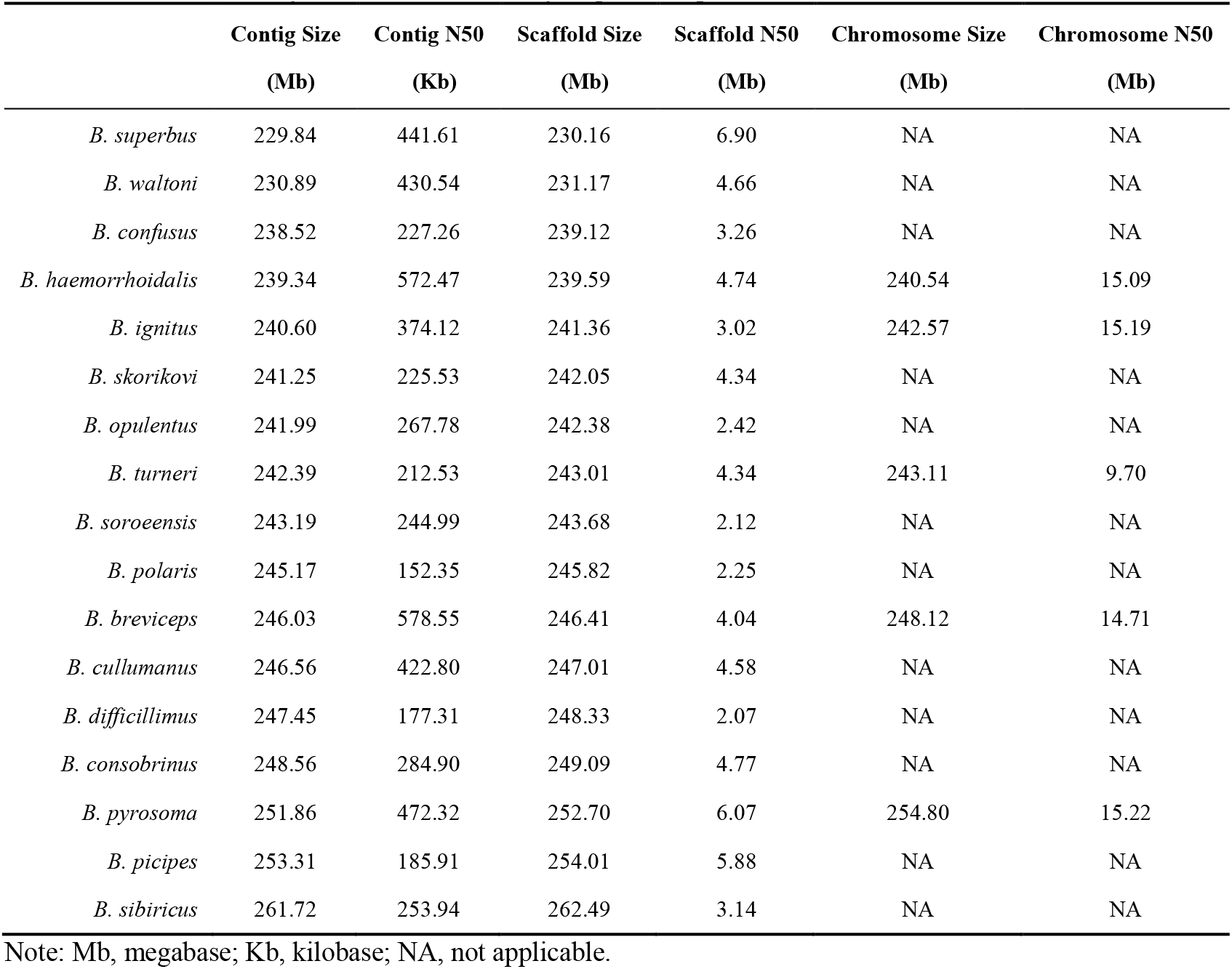
Genome assembly results of the 17 newly sequenced species.

|  | Contig Size<br>(Mb) | Contig N50<br>(Kb) | Scaffold Size<br>(Mb) | Scaffold N50<br>(Mb) | Chromosome Size<br>(Mb) | Chromosome N50<br>(Mb) |
| --- | --- | --- | --- | --- | --- | --- |
| *B. superbus* | 229.84 | 441.61 | 230.16 | 6.90 | NA | NA |
| *B. waltoni* | 230.89 | 430.54 | 231.17 | 4.66 | NA | NA |
| *B. confusus* | 238.52 | 227.26 | 239.12 | 3.26 | NA | NA |
| *B. haemorrhoidalis* | 239.34 | 572.47 | 239.59 | 4.74 | 240.54 | 15.09 |
| *B. ignitus* | 240.60 | 374.12 | 241.36 | 3.02 | 242.57 | 15.19 |
| *B. skorikovi* | 241.25 | 225.53 | 242.05 | 4.34 | NA | NA |
| *B. opulentus* | 241.99 | 267.78 | 242.38 | 2.42 | NA | NA |
| *B. turneri* | 242.39 | 212.53 | 243.01 | 4.34 | 243.11 | 9.70 |
| *B. soroensis* | 243.19 | 244.99 | 243.68 | 2.12 | NA | NA |
| *B. polaris* | 245.17 | 152.35 | 245.82 | 2.25 | NA | NA |
| *B. breviceps* | 246.03 | 578.55 | 246.41 | 4.04 | 248.12 | 14.71 |
| *B. cullumanus* | 246.56 | 422.80 | 247.01 | 4.58 | NA | NA |
| *B. difficillimus* | 247.45 | 177.31 | 248.33 | 2.07 | NA | NA |
| *B. consobrinus* | 248.56 | 284.90 | 249.09 | 4.77 | NA | NA |
| *B. pyrosoma* | 251.86 | 472.32 | 252.70 | 6.07 | 254.80 | 15.22 |
| *B. picipes* | 253.31 | 185.91 | 254.01 | 5.88 | NA | NA |
| *B. sibiricus* | 261.72 | 253.94 | 262.49 | 3.14 | NA | NA |

Note: Mb, megabase; Kb, kilobase; NA, not applicable.

Sequencing and assembly strategies included generating two Illumina sequencing datasets for each species: (i) overlapping paired-end reads (2 × 250 bp) from one small-insert fragment library (insert size: 400 or 450 bp); and (ii) paired-end reads (2 × 150 bp) from four large-insert jump libraries (insert sizes: 4 kb, 6kb, 8kb and 10 kb, respectively; Additional file 1: Table S2). Whole-genome overlapping paired-end reads from fragment libraries were assembled into continuous sequences (contigs) using the software DISCOVAR *de novo* (Love, et al. 2016), then scaffolded with reads from jump libraries using the software BESST (Sahlin, et al. 2014). The resulting assemblies have a mean contig N50 of 325 Kb, ranging up to 579 Kb for *B. breviceps;* the mean scaffold N50 is 4.0 Mb, ranging up to 6.9 Mb for *B. superbus* (Table 1). Genome assembly quality in terms of expected gene content was evaluated by Benchmarking Universal Single-Copy Ortholog (BUSCO) analysis (Waterhouse, et al. 2018), which showed high BUSCO completeness scores (average 99.0%, from 97.5 to 99.6%; Additional file 2: Figure S1) for all genomes.

Genome annotation resulted in total protein-coding gene predictions per species ranging from 14,027–16,970 (mean = 15,838, standard deviation = 908; Additional file 1: Table S3). These were annotated using the MAKER pipeline (Cantarel, et al. 2008), based on *ab initio* gene predictions, transcript evidence, and homologous protein evidence. Gene counts are similar to those of 12 drosophilid species (mean = 15,361, sd. = 852 Clark et al., 2007), but are higher than those of 19 anophelines (mean = 13,110, sd. = 1,397) (Neafsey, et al. 2015), and they do not correlate with assembly contiguity (p = 0.1757; Additional file 2: Figure S2). Between 7,299–8,135 genes were assigned at least one Gene Ontology (GO) term and 9,431–10,578 genes were annotated with at least one protein domain (Additional file 1: Table S3). BUSCO analysis of the annotated genes also showed high completeness scores for all species (Additional file 2: Figure S3). Furthermore, comprehensive miRNA, tRNA, and lncRNA gene prediction revealed an average of 93, 306, and 3,353 genes, respectively (Additional file 1: Table S3). Finally, transposable element (TE) annotation showed that the total TE content ranged from 9.66% (22.2 Mb) in *B. superbus* to 17.88% (46.9 Mb) in *B. sibiricus* (Additional file 1: Table S4).

### Genome-scale phylogeny of bumblebees

The species-level molecular phylogeny (Figure 1A) estimated from maximumlikelihood analysis with IQTree (Minh, et al. 2020b) is largely consistent with previously inferred phylogenetic relationships of the 15 subgenera based on five genes (Cameron, et al. 2007; Williams, et al. 2008), showing only two topological differences. The results support previous conclusions that: (i) subgenus *Mendacibobus* (*Md*) is the sister group to all the other subgenera; and (ii) lineages named *Psithyrus (Ps)* are within the *Bombus* clade, arguing they should not be named as an independent genus (Figure 1A). The species phylogeny was built from the concatenated aligned protein sequences of 3,617 universal single-copy orthologs from 19 bumblebee species (17 from the current study, two published previously: *B. terrestris* and *B. impatiens* (Sadd, et al. 2015)) and four honeybee species *(A. florea, A. dorsata* (Oppenheim, et al. 2020), *A. cerana* (Park, et al. 2015), and *A. mellifera* (Weinstock, et al. 2006)), with orthologous groups delineated using the OrthoDB software (Kriventseva, et al. 2015). Complementary analysis with ASTRAL (Zhang, et al. 2018) resulted in an identical species tree with the exception of the placement of *B. pyrosoma,* which no longer forms a monophyletic pairing with *B. breviceps* (Additional file 2: Figure S4). This type of discordance between species tree methods is consistent with a known shortcoming of maximum-likelihood concatenation in the presence of incomplete lineage sorting (ILS) (Kubatko and Degnan 2007; Mendes and Hahn 2018), implying that the ASTRAL topology is likely the correct topology.

**Figure 1.**
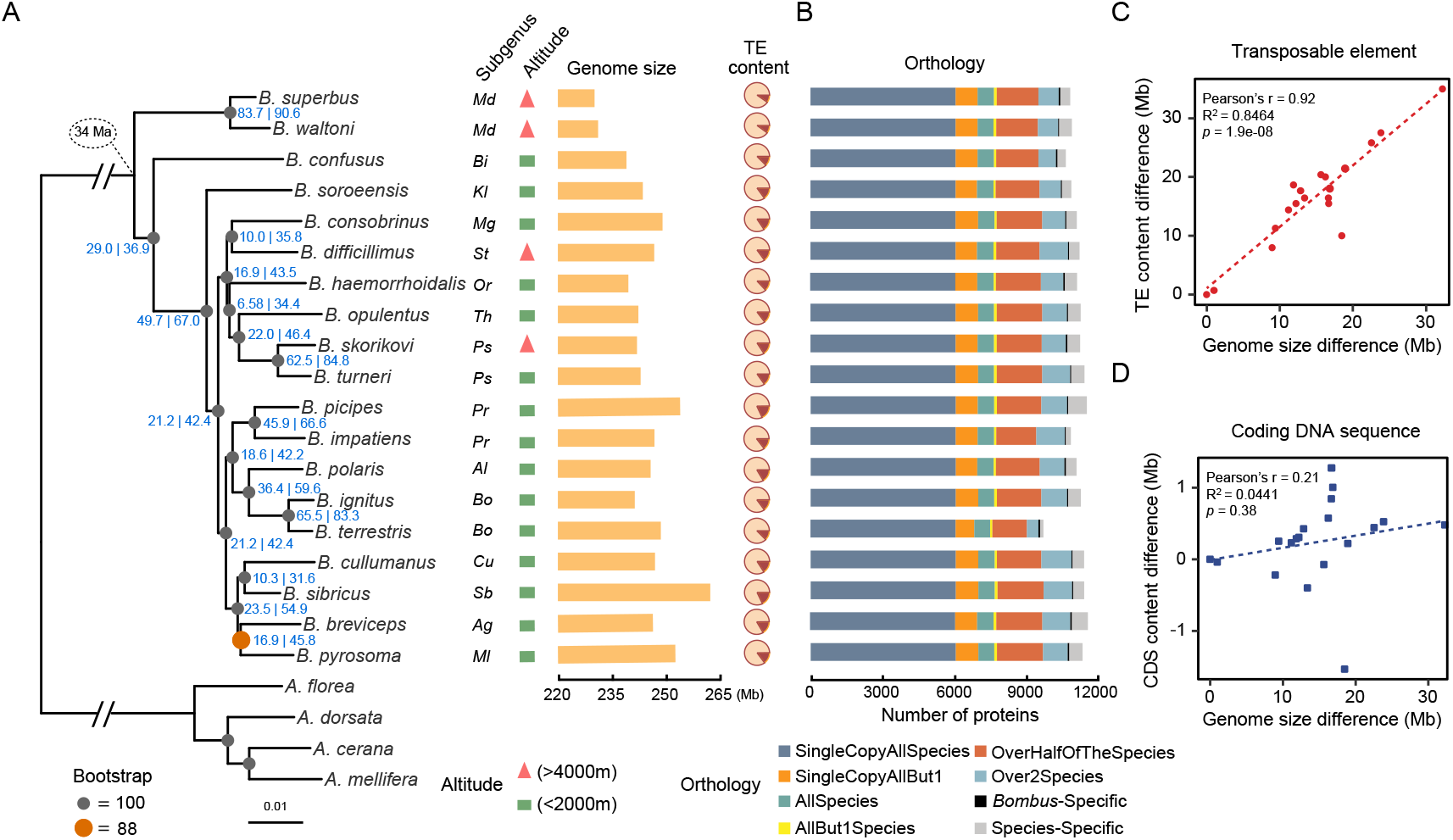
Phylogenetic, genomic and proteomic comparisons of 19 bumblebee species. (**A**) From left to right: the maximum likelihood molecular species phylogeny built from 3,617 concatenated single-copy orthologous groups from all sequenced bumblebees and honeybee outgroups. Node labels in blue are of the following format: gene concordance factors | site concordance factors. Branches scaled by relative number of substitutions; the subgenus that each bumblebee species belongs to (Md, *Mendacibombus;* Bi, *Bombias;* Kl, *Kallobombus;* Mg, *Megabombus;* St, *Subterraneobombus;* Or, *Orientalibombus;* Th, *Thoracobombus;* Ps, *Psithyrus;* Cu, *Cullumanobombus;* Sb, *Sibiricobombus*; Ag, *Alpigenobombus*; Ml, *Melanobombus*; Pr, *Pyrobombus*; Al, *Alpinobombus*; Bo, *Bombus*); altitude of species collection site (red triangle: extreme high-altitude; green rectangle: low-altitude); genome assembly size of each sequenced species; fraction of transposable elements (TE) (brown) in each genome. (**B**) Bar plots show total gene counts for each bumblebee partitioned according to their orthology profiles, from ancient genes found across bumblebees to lineage-restricted and species-specific genes. (**C**) and (**D**) represent the contribution of transposable element and coding DNA sequence to genome size variation across bumblebees, respectively. Differences in the total content of transposable elements (**C**) and coding DNA sequences (**D**) of the 19 genomes relative to that of *B. superbus* (which has the smallest genome assembly size) are plotted against their genome size differences (relative to that of *B. superbus*).

However, inferring rooted gene trees from 3,530 single-copy orthologous groups reveals extreme levels of discordance: none of their topologies match the topology of the tree inferred from concatenation (Additional file 1: Table S5 and Additional file 1: Table S6), and nearly every gene tree has a unique topology (Additional file 1: Table S7). Such extreme levels of discordance have been observed previously in birds (Jarvis, et al. 2014) and tomatoes (Pease, et al. 2016), and have been attributed to a variety of sources, such as ILS and introgression (Maddison 1997). A lack of informative sites, only 24%, compared to 47% in a similar dataset of 25 drosophilids (Da Lage, et al. 2019), possibly due to the relatively recent diversification of bumblebees (Hines 2008), may also cause discordance. Concordance analysis (Minh, et al. 2018) shows that, on average, nodes in the species tree are present in only a third of gene trees and only about half of informative sites support the species tree (node labels in Figure 1A). These site concordance factors, the short internal branches of the species tree, and the strong correlation between them (Additional file 2: Figure S5), are consistent with ILS driving the observed gene tree discordance. Gene-level phylogenies are therefore used in all subsequent gene-based molecular evolution analyses because such discordance can bias inferences of substitutions when mapped onto a species tree (Mendes and Hahn 2016).

### Major genomic rearrangements in social parasites

The five Hi-C genome assemblies indicate that four of the five subgenera have 18 chromosomes (Figure 2A and 2C; Additional file 2: Figure S6A-B), consistent with previous karyotypic analysis that inferred the ancestral chromosome number is 18 (Owen, et al. 1995). However, the social parasite bumblebee, *B. turneri,* subgenus *Psithyrus*, has 25 chromosomes (Figure 2B), consistent with previous cytological work (Owen and Robin 1983). Despite the higher chromosome number, its genome size is within the range of other bumblebees (Figure 1; Table 1). Pairwise comparisons between *B. turneri* and each of the other four chromosomal-level assemblies to investigate macrosynteny relationships and understand how a 25-chromosome karyotype was derived from the ancestral state revealed three processes. First, some chromosomes descended, structurally unchanged, from ancestral chromosomes (e.g., chromosome 5; Figure 2D in blue). Second, some originated by fission of an ancestral chromosome (e.g., 11 and 25 of *B. turneri* originated by the fission of ancestral chromosome 11; Figure 2D in red). Lastly, some are derived from fusions of two or more ancestral chromosome segments (e.g., *B. turneri* chromosome 22 was derived from the fusion of segments of ancestral chromosomes 7, 8, 10, and 16 (Figure 2D in gold). Pairwise comparisons between *Psithyrus* and members of other subgenera reveal similar results, and support the inference that the 25 chromosomes of the social parasite bumblebee result from a combination of fission, fusion, and retention of ancestral chromosomes (Additional file 2: Figure S6).

**Figure 2.**
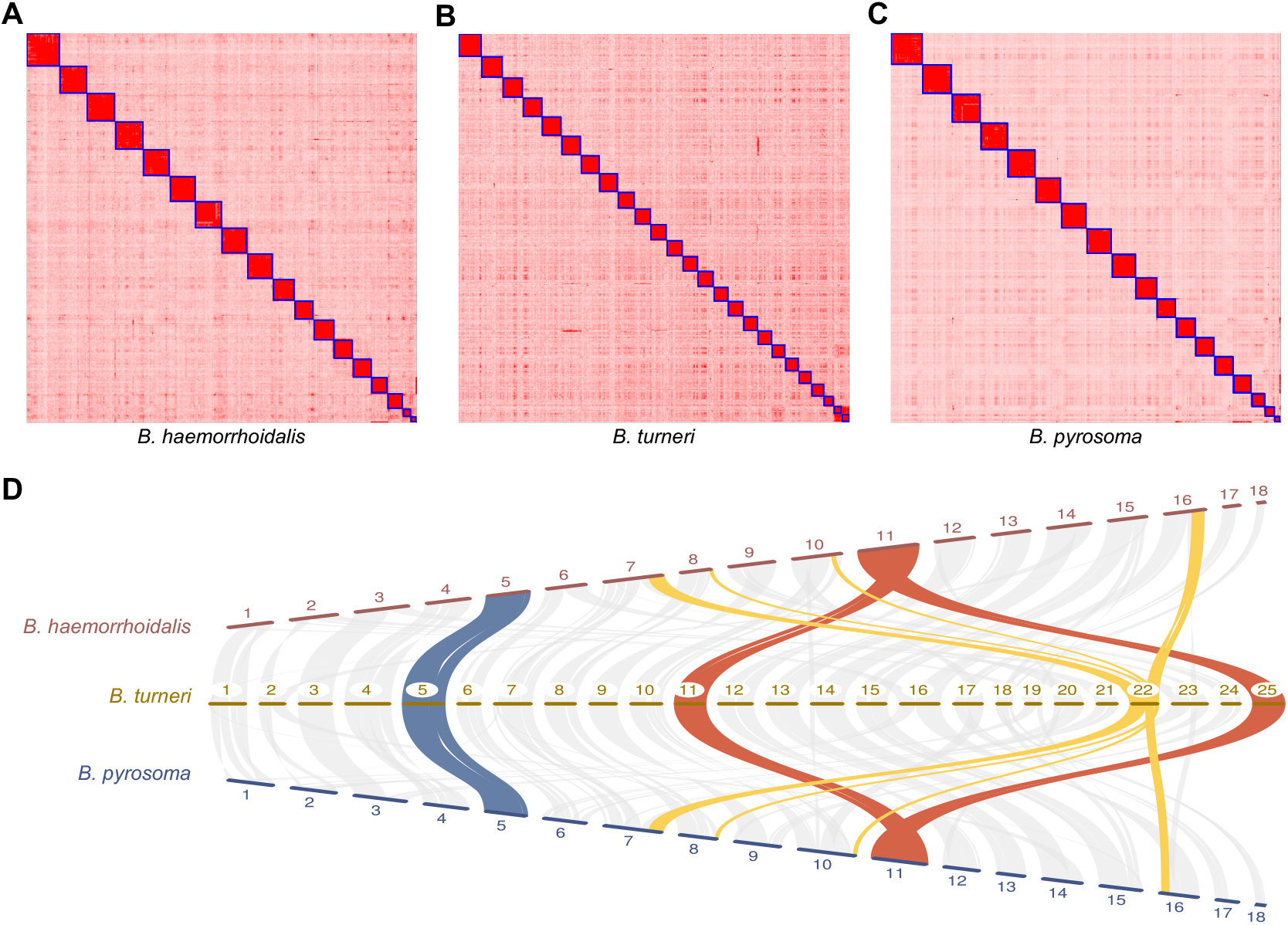
Chromosome number evolution in representative *Bombus* species. Hi-C contact heatmaps for *B. haemorrhoidalis* (**A**), *B. turneri* (**B**), and *B. pyrosoma* (**C**) show that the three species have 18, 25, and 18 chromosomes, respectively. The 18-chromosome karyotype is the inferred ancestral genome structure, with 25 chromosomes found in social parasite bumblebees of the subgenus *Psithyrus.* (**D**) Macrosynteny comparisons across *B. haemorrhoidalis*, *B. turneri* and *B. pyrosoma* shows how the 25 *B. turneri* chromosomes result from a combination of fission (red), fusion (yellow), and retention (blue) of ancestral chromosomes.

Rates of chromosome evolution, in terms of rearrangements relative to *B. terrestris,* were investigated for each of the five species with chromosome-level assemblies. Rearrangement rates in bumblebees range from 0.0016–0.0075 inversions/Mb/My (Additional file 1: Table S8), which is much lower than those of drosophilids (0.013–0.159 inversions/Mb/My) and anophelines (0.052–0.068 inversions/Mb/My) (Neafsey, et al. 2015; von Grotthuss, et al. 2010). Thus, although bumblebee genomes have a high recombination rate (Wilfert, et al. 2007), their rates of chromosome evolution are relatively slow, which is further supported by the observed high synteny contiguity across species (average 88%, from 80–95%; Additional file 1: Table S9).

### Transposable elements drive genome size variation

Genome assembly sizes (haploid) range from 230 Mb in *B. superbus* to 262 Mb in *B. sibiricus* (Figure 1). Ancestral genome size inference of bumblebees produced an estimate of 230-231 Mb, similar to that of members of the subgenus *Mendacibombus*, but smaller than the genomes of all other extant bumblebees surveyed (Additional file 2: Figure S7).

Comparing genome size differences with relative content of TEs, simple sequence repeats (SSRs), and coding DNA sequences (CDS) shows that TE content explains a majority of the differences across bumblebees (Pearson correlation R = 0.92, *P* = 1.9e-08, R^2^=0.85; Figure 1C, Figure 1D, Additional file 2: Figure S8). *Mendacibombus* species have a smaller genome size than other species (Figure 1), and TEs that transposed in non-*Mendacibombus* species after divergence from *Mendacibombus* show copy numbers ranging from 1,992–4,755 (Additional file 2: Figure S9), supporting the contribution of TEs to genome size evolution. Furthermore, TE proliferation history analysis indicated that all non-*Mendacibombus* species have more recent TE amplification peaks (Additional file 2: Figure S10), consistent with increased TE activity driving genome size increases.

The genomic distributions of TEs include 1,074–1,786 TE loci that overlap with the coding regions of protein-coding genes (Additional file 1: Table S10), and 352 of these genes are universal single-copy across the 19 bumblebees whose *dN/dS* values are all < 1 (Additional file 1: Table S11), indicating that TEs may have been exonized in bumblebee genomes to form novel proteins. In addition, there are thousands of TEs located within 1 kb of a gene in each species (Additional file 1: Table S10), and, in *B. terrestris*, 278 such TEs co-locate with open chromatin regions detected by ATAC-seq (Additional file 1: Table S12), suggesting those TEs may have become incorporated into regulatory sequences.

### Gene content evolution reflects foraging and diet diversity

Orthology delineation results indicate that a majority of genes are found in one or more copies in nearly all lineages across bumblebees (Figure 1B). These include 53 *Bombus*-specific ortholog groups, which are present in all 19 bumblebees but absent in all four honeybees (Figure 1B; Additional file 1: Table S13), and may play roles in lineage-specific traits. Functional annotation suggests that five of these *Bombus*-specific genes are associated with protein metabolism and transport (Additional file 1: Table S13), potentially linked to the higher protein content of pollen collected by bumblebees than honeybees (Leonhardt and Blüthgen 2011). Ortholog groups with the broadest species representation are functionally enriched for core biological processes such as protein transport, signal transduction (e.g. Wnt pathway), (de)ubiquitination, and cytoskeleton organization (Additional file 1: Table S14). In contrast, those with sparse or lineage-restricted species representation are enriched for processes including smell and taste perception, amino acid biosynthesis, and oxidation-reduction (Additional file 1: Table S14). On average, 465 species-specific genes (those without an ortholog in any other lineage) were identified in each bumblebee species (range 137–767) (Additional file 1: Table S15), which may contribute to species-specific traits but whose functional roles remain to be explored.

Turnover analysis of gene repertoires across the *Bombus* phylogeny (15 species, one per subgenus) using CAFE v3.0 (Han, et al. 2013) identified expansions and contractions among 13,828 gene families and quantified variations in gene gain/loss rates across species (Additional file 2: Figure S11). After error correction, the overall rate of gene gain/loss in *Bombus* genomes is 0.0036/gene/million years, similar to an analysis of 18 anopheline species and 25 drosophilids (Additional file 1: Table S16) (Da Lage, et al. 2019; Neafsey, et al. 2015). However, these genus-specific gene gain/loss rates are 2-3 times higher than order-wide rates, which average 0.0011 (Additional file 1: Table S16) (Thomas, et al. 2020), possibly due to the denser sampling in genus-level studies that allow more events to be captured. Gene gain and loss events, along with the number of rapidly evolving gene families, are summarized for each species (Additional file 1: Table S17), with a total of 3,797 rapidly changing gene families. The most dynamic gene families are enriched for processes including smell and taste perception, chitin metabolism, microtubule-based movement, and methylation (Additional file 1: Table S18). Complementary analysis using three measures of gene copy number variation also identifies these processes as enriched among the most variable gene families, in contrast to the most stable that are involved in processes related to translation, adhesion, and transport (Additional file 1: Table S19). In terms of protein domain copy number evolution, the most highly variable genes are those with protein-protein interaction mediating F-box domains, putatively DNA-binding SAP motifs, and phosphate-transferring guanylate kinases (Additional file 1: Table S20).

### Stable intron-exon structures with abundant stop-codon readthrough

Protein-coding potential analysis using *B. terrestris* as the reference species identified 851 candidate readthrough stop codons (Additional file 2: Figure S12; Additional file 1: Table S21), i.e. where translation likely continues through stop codons to produce extended protein isoforms. Coding potential was assessed using PhyloCSF (Lin, et al. 2011) on whole genome alignments of all 19 bumblebees and four honeybees. The false discovery rate was estimated using enrichment for the TGA-C stop codon context, which is favored in readthrough genes, to infer that no more than 30% of the 200 highest-scoring candidates are false positives, and that at least 306 of our 851 candidates undergo functional readthrough. While rare beyond Pancrustacea, hundreds of *Drosophila* and *Anopheles* genes undergo readthrough, and in Hymenoptera estimates for honeybee are low but for *Nasonia* wasps high (Dunn, et al. 2013; Jungreis, et al. 2016; Jungreis, et al. 2011; Rajput, et al. 2019). These whole-genome-alignment-based results support the prediction (Jungreis, et al. 2011) that insect species have abundant stop-codon readthrough.

In contrast, intron-exon boundaries within *Bombus* genes are relatively stable. Examining evolutionary histories of intron gains and losses revealed few changes, representing only 3-4% of ancestral intron sites, with more gains than losses (Additional file 2: Figure S13; Additional file 1: Table S22), unlike drosophilids and anophelines where losses dominate (Neafsey, et al. 2015), suggesting that bumblebee gene structure has remained relatively stable over the 34 million years since their last common ancestor.

### Divergence and selective constraints of protein-coding genes

Bumblebee genes with elevated sequence divergence and/or relaxed constraints include processes related to smell perception, chitin metabolism, RNA processing, DNA repair, and oxidation-reduction (Figure 3). Measures of evolutionary rate (amino acid sequence divergence) and selective constraint (*dN/dS*) showed similar trends among different functional categories of genes. Most genes are strongly constrained, with median estimates of *dN/dS* much lower than one. Assignment of GO terms and InterPro domains is usually biased towards slower-evolving, well-conserved genes (Additional file 2: Figure S14). Nevertheless, functional categories with the fastest-evolving genes are further supported and complemented by examining molecular function GO terms (Additional file 2: Figure S15A) and InterPro domains (Additional file 2: Figure S15B), which show elevated rates for odorant binding, olfactory receptor activity, chitin binding, oxidoreductase activity, serine-type endopeptidase activity, and olfactory receptor domains. GO term enrichment analysis of the slowest and fastest evolving subsets of genes, bottom and top 20% respectively (Additional file 2: Figure S16), showed genes with the slowest evolutionary rates and the lowest *dN/dS* ratios were enriched for essential housekeeping biological processes and molecular functions (Additional file 1: Table S23; Additional file 1: Table S24). In contrast, genes with the fastest evolutionary rates were enriched for processes linked to polysaccharide biosynthesis, tRNA aminoacylation, drug binding and RNA methyltransferase activity (Additional file 1: Table S23). Genes with the highest *dN/dS* ratios were enriched for processes and functions including proteolysis, translation, ncRNA processing, and chitin metabolism (Additional file 1: Table S24).

**Figure 3.**
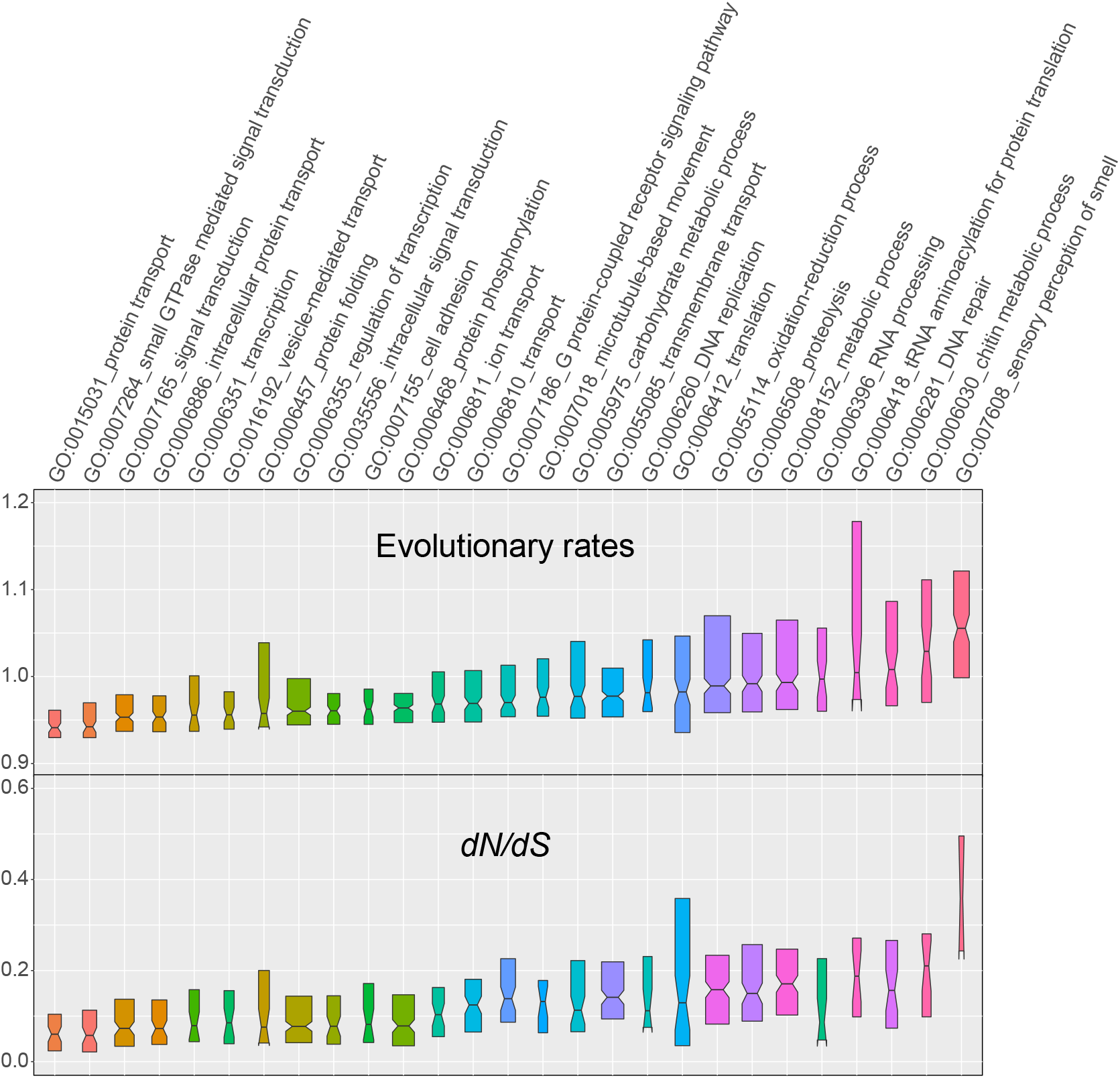
Molecular evolution of protein-coding genes in terms of evolutionary rate (amino acid sequence divergence) and *dN/dS* ratio among selected Gene Ontology (GO) Biological Process terms. Categories are sorted by evolutionary rate from the most conservative (left) to the most dynamic (right) and colored from the highest values (red) to the median value (blue) to the lowest values (orange). Notched boxes show medians of orthologous group values with the limits of the upper and lower quartiles, and box widths are proportional to the number of orthologous groups in each category.

### Codon usage bias driven by AT content

Analysis of codon usage bias showed no evidence for selection on optimal codons, in contrast to drosophilids but similar to anophelines (Neafsey, et al. 2015; Vicario, et al. 2007). Instead, codon usage bias in bumblebees seems to be driven mainly by AT content, consistent with previous reports in Hymenoptera (Behura and Severson 2012). Optimal codons were estimated in each species and correlation coefficients were computed between relative synonymous codon usage (RSCU) and effective number of codons (ENC) per gene. All species have a similar preference and intensity of preference; for each amino acid, there was a consistently highly preferred codon and often a secondarily preferred one, all ending in A/T (Additional file 2: Figure S17). To test if codon usage could largely be explained by mutation bias, a linear model was used to predict Fop (frequency of optimal codon) from overall gene AT content and amino acid use. The model explained 99.2% of the Fop variation without the need to include the species origin of each gene. The AT content alone explained 81% of the variation (Additional file 2: Figure S18). Moreover, a strong correlation was observed between codon AT content and the correlation between RSCU and ENC across all species (Additional file 2: Figure S19).

### Evolution of genes associated with bumblebee eco-ethology

Many ecological and environmental factors—for example, shortage of food, pathogen emergence, pesticide exposure, and climate change—are contributing to the overall decline of bumblebees worldwide (Cameron and Sadd 2019; Goulson, et al. 2015; Williams, et al. 2009). To begin to explore the complement of genes likely to be involved in bumblebee interactions with their environment, we examined the evolution of gene families associated with their ecology and life histories. Sampling across the *Bombus* genus enabled the first survey of natural gene repertoire diversity of such families that are likely to be important for bumblebee adaptability and success.

#### Chemosensory receptor diversity

Chemosensation plays a critical role in locating food and nests, communicating with nestmates, and identifying other environmental cues (Ayasse and Jarau 2014). A search of the three major chemosensory receptor gene families—odorant receptors (ORs), gustatory receptors (GRs), and ionotropic receptors (IRs)—in the sequenced bumblebee genomes identified 3,228 genes (Additional file 1: Table S25). Only complete genes were used for gene gain and loss analysis. Despite the similarities in total OR gene counts, examples of gene gain/loss were observed in specific lineages. There was a net loss of 15 ORs in the common ancestor of the subgenus *Mendacibombus (Md)* (Figure 4A; Additional file 2: Figure S20). Species in *Mendacibombus* mainly inhabit high mountains including the Qinghai-Tibetan plateau, with relatively low floral diversity (Williams, et al. 2018), which may be linked to OR loss in this lineage. A net loss of 11 ORs was observed in the common ancestor of subgenus *Psithyrus (Ps)* (Figure 4A; Additional file 2: Figure S20). For ORs shared across bumblebees, eight showed evidence of positive selection in a subset of species, including putative pheromone receptors (Additional file 1: Table S26). Compared with ORs, GRs and IRs have much lower and more stable gene counts (Additional file 2: Figure S20). However, despite overall conservation of gene number and widespread evidence for purifying selection, there is evidence that some GR and IR genes experienced positive selection in a subset of species, including receptors putatively involved in sensing fructose and temperature (Additional file 1: Table S26).

**Figure 4.**
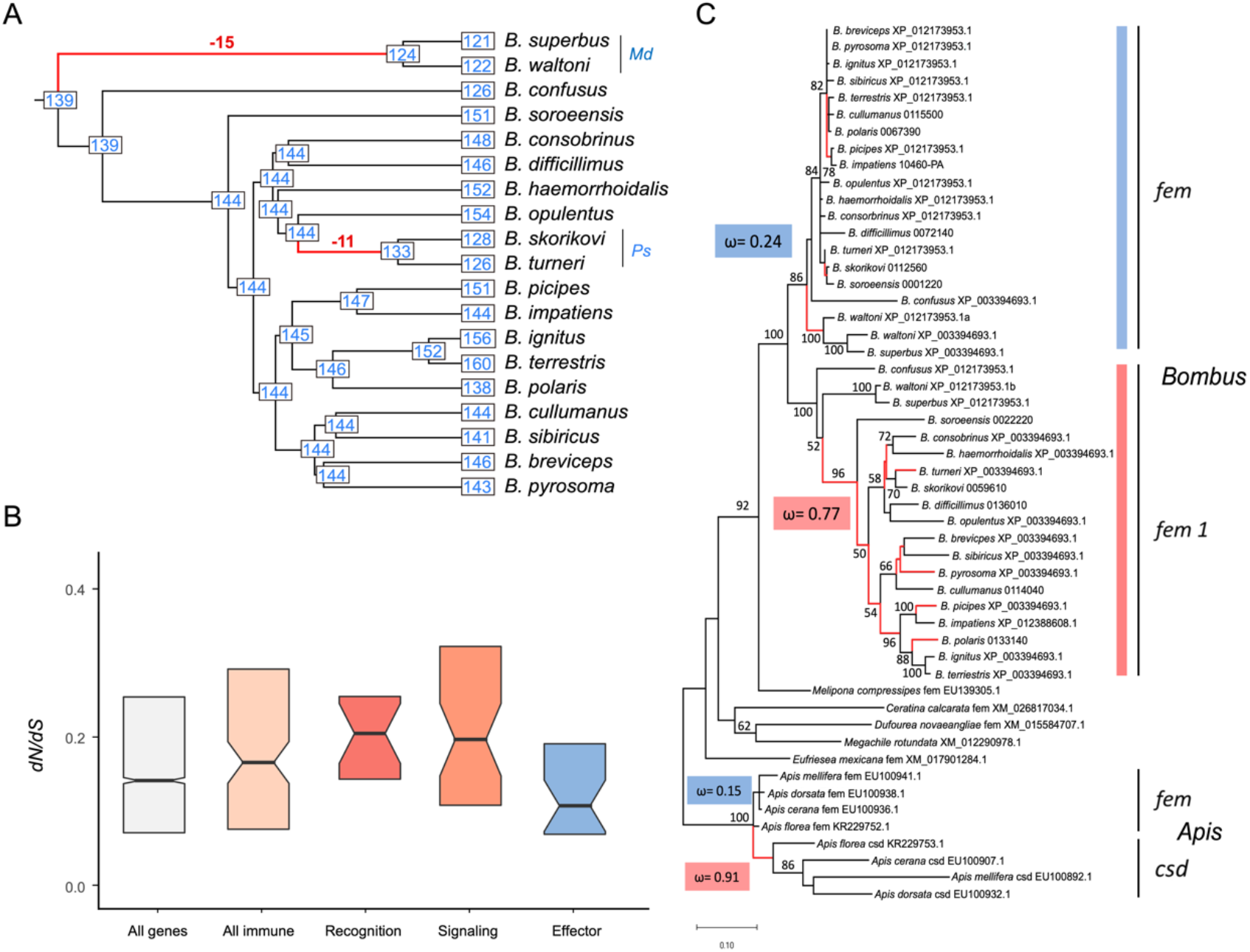
Evolution of genes associated with ecology and reproduction. (**A**). Observed gene counts and inferred ancestral gene counts of bumblebee odorant receptors (ORs) on an ultrametric phylogeny, highlighting two major gene loss events (the complete result is available in Additional file 2: Figure S21). *Md*, *Mendacibombus*; *Ps*, *Psithyrus.* (**B**). Boxplots showing *dN/dS* ratios for different categories of immune genes and all single-copy genes in bumblebee (All genes). Elevated *dN/dS* ratios among immune-related genes is driven by higher ratios for genes involved in recognition and signaling processes. Notched boxes show medians of orthologous group values with the limits of the upper and lower quartiles. (C). The evolutionary history of fem genes of bees including their paralogs *fem1* in *Bombus* and *csd* in *Apis*. Global non-synonymous to synonymous rate ratio (ω) were calculated for *femBombus* (reference, blue) and *fem1 Bombus* (test, red), including a branch-site testing framework with model fitting and Likelihood Ratio Tests, showing evidence for relaxation of selection in*fem1 Bombus* (P<0.001, LR = 36.34). Spurious actions of diversifying selection on branches predominantly found in *fem1 Bombus* are marked in red. For comparison, ω for fem and *csd* in *Apis* is given, known as striking example of neofunctionalization.

#### Detoxification capacity

Detoxification genes are used to neutralize toxic plant secondary metabolites and pesticides. Repertoires of carboxyl/cholinesterases (CCEs), cytochrome P450 monooxygenases (P450s), and glutathione S-transferases (GSTs) in the 17 genomes are much smaller than in drosophilids and anophelines (Additional file 1: Table S27), indicating a genus-wide deficit of this gene category, previously observed in two bumblebees (Sadd, et al. 2015). There are 88 detoxification genes on average in bumblebees, with little variation across species (Additional file 1: Table S27). Despite overall conservation of gene number and widespread evidence for purifying selection (mean *dN/dS* is 0.26), a total of 30 detoxification genes, including CCEs, P450s, and GSTs, showed evidence of positive diversifying selection in a subset of species (Additional file 1: Table S28).

#### Immune defense

Immune genes are involved in recognition of and defense against pathogens. Similar to detoxification genes, counts in the 17 sequenced genomes are much lower than in drosophilids and anophelines (Additional file 1: Table S29), showing that the previously noted paucity in two bumblebees (Barribeau, et al. 2015; Sadd, et al. 2015) extends to the whole genus. Bumblebee genomes contain components of all major immune pathways described in insects, and gene counts are fairly conserved across species (Additional file 1: Table S29). For example, all species have two genes encoding Gram-negative bacteria binding-proteins, while peptidoglycan-recognition proteins are more variable with between four and six gene copies. Comparing *dN/dS* ratios between immune genes and all single-copy orthologous genes in bumblebees showed that immune genes exhibit slightly higher *dN/dS* ratios (P = 0.04, Wilcoxon rank sum test), and among immune genes, recognition and signaling genes have higher *dN/dS* ratios than effector genes (Figure 4B). In addition, despite widespread evidence for purifying selection, a total of 77 immune genes showed evidence of positive selection in a subset of bumblebee species (Additional file 1: Table S30). *B. cullumanus, B. difficillimus,* and *B. confusus,* which have no reported internal parasites (Arbetman, et al. 2017), are among the species that have the most positively selected genes (Additional file 1: Table S30).

### Genes involved in high-elevation adaptation

*Bombus superbus, B. waltoni, B. difficillimus,* and *B. skorikovi* are four species collected at elevations > 4,000 m that represent three subgenera (Figure 1). No genes show signatures of positive selection in all high-elevation species but none of the low-elevation species. However, nine genes show evidence of positive selection in species representing two of the three high-elevation subgenera, but none of the low-elevation species (Additional file 1: Table S31). Two encode Myosin-VIIa and CPAMD8, respectively, which are involved in eye development (Cheong, et al. 2016; Williams and Lopes 2011). As bumblebees detect flowers visually (Meyer-Rochow 2019), signatures of selection might be related to fine tuning eye development for optimal foraging in high altitude light conditions. Three genes encode histone deacetylase, synaptotagmin-12, and heterogeneous nuclear ribonucleoprotein, which are involved in maintaining muscle integrity and keeping “flight state”, which is critical for undertaking long-distance food-searching (Liu, et al. 2001; Manjila, et al. 2019; Pigna, et al. 2019). Three genes encode sodium-coupled monocarboxylate transporter, glycosyltransferase family protein, and xyloside xylosyltransferase 1, these genes are believed to be involved in metabolic adaptation to hypoxia (Gustafsson, et al. 2005; Lee, et al. 2013; Shirato, et al. 2010; Véga, et al. 2006) (Additional file 1: Table S31). The remaining gene encodes a proton channel, which may be also involved in the metabolic adaptation to hypoxia (Bacon and Harris 2004).

#### Sex-determination

Evolutionary analysis of sex-determination genes in bumblebees and related species indicated that all *Bombus* genomes share a duplicated copy of *feminizer (fem),* named *fem 1* (Figure 4C). Compared to *fem, fem 1* shows a higher level of divergence among bumblebees (*fem*_Bombus_ *dN/dS* = 0.24; *fem 1*_Bombus_ *dN/dS* = 0.77; Figure 4C). These ratios are close to the range observed for *Apis,* in which *fem* has evolved under purifying selection and the paralogous gene *complementary sex determiner* (*csd*) has evolved by neo-functionalization (Figure 4C) (Hasselmann, et al. 2008). A hypothesis branch-site testing framework (RELAX), identifies evidence for relaxation of selection in *fem 1_Bombus_* compared to *fem_Bombus_* (P<0.001, LR = 36.34). Moreover, the spurious action of diversifying selection on branches was predominantly found in *fem 1_Bombus_* (Figure 4C). A mixed effect model of evolution (MEME) was applied to identify individual sites that were subject to episodic diversifying selection, and at least 15 sites (p< 0.05) were found to be under positive selection, with some being located in known motifs (Additional file 2: Figure S21). The results of these selection analyses suggest that both *fem* and *fem 1* contribute to the *Bombus* sex determination pathway. For the *transformer 2 (tra-2)* gene, consistent amino acid changes between *Bombus* and *Apis* were found within the RNA recognition domain (Additional file 2: Figure S22), supporting a previous hypothesis of a regulatory modification between the two groups (Biewer, et al. 2015).

## Discussion

Comparative analysis of multiple genomes in a phylogenetic framework substantially improves the precision and sensitivity of evolutionary inference and provides robust results identifying stable and dynamic features. In this study, we performed comparative analyses of genome structures and contents, as well as global and family-targeted gene evolutionary dynamics across the phylogeny of *Bombus,* using 17 annotated *de novo* assemblies and two previously published genomes.

Many attributes of bumblebee genomes are highly conserved across species. For example, overall genome size and genome structure, the number of protein-coding genes and non-coding RNAs, gene intron-exon structures, and the pattern of codon usage are all very similar across these 19 genomes. However, other aspects of genome biology are dynamically evolving. TEs are a major contributor to genome size variation (Figure 1) as well as a potential source of coding and regulatory sequences (Additional file 1: Table S10-12). Differential gene gain and loss also contribute to gene content variation across bumblebees and lead to lineage-specific gene repertoires (Figure 4 A; Additional file 2: Figure S20; Additional file 1: Table S17). Finally, for genes shared by all species, the action of positive selection is different across species (Additional file 1: Table S26; Additional file 1: Table S28; Additional file 1: Table S30; Additional file 1: Table S31), which can lead to gene functional divergence possibly reflecting key eco-ethological differences.

An exception to the otherwise overall conserved genome structure is the set of species in the subgenus *Psithyrus*. These bumblebees exhibit social parasitism; they do not have a worker caste, and it is not necessary for them to forage for nectar and pollen (Lhomme and Hines 2019). Originally, this subgenus was argued to be a separate genus due to distinct behavior and higher chromosome number, however subsequent phylogenetic analysis placed *Psithyrus* within the subgenus *Bombus* (Williams, et al. 2008). Here, based on a much larger genomic dataset, we confirm that species in the subgenus form a monophyletic group within the *Bombus* clade (Figure 1A). In addition, we show that, although *Psithyrus* species have an increased chromosome number, their genome sizes are within the range of those of the other bumblebees (Figure 1A), and their 25 chromosomes reflect a mix of fission, fusion, and retention of the 18 ancestral bumblebee chromosomes (Figure 2; Additional file 2: Figure S6). Chromosome rearrangements (e.g., fissions, fusions, and inversions) have been posited to play roles in speciation (Ayala and Coluzzi 2005), and thus may explain the diversification and social parasitic behavior of *Psithyrus*. In addition to genome structure variation, we identified a net loss of 11 odorant receptor genes in the common ancestor of *Psithyrus* species (Figure 4), which could be a cause or consequence of their socially parasitic behavior.

Bumblebee species exhibit different food preferences (Goulson and Darvill 2004; Sikora and Kelm 2012; Somme, et al. 2015), but the genetic basis underlying such variation is unknown. Like in other insects, smell and taste are used to distinguish different food sources (Kunze and Gumbert 2001; Ruedenauer, et al. 2015). In this study, we found out that genes involved in smell and taste perception are among the fastest evolving gene categories, both in copy number variation and in sequence divergence (Figure 3; Additional file 2: Figure S15; Additional file 1: Table S18-19). Therefore, the dynamic evolution of genes involved in smell and taste perception likely contribute to different food preferences, improved understanding of which could inform the use of new species in agricultural settings. Bumblebees exhibit rich morphology differences across species (Williams 1994) and they show species-specific responses to insecticides (Baron, et al. 2017). Chitin is a major component of the insect cuticle and peritrophic matrix, and chitin metabolic processes are related to morphogenesis, resistance to insecticides, and the tolerance of toxins in food (Barbehenn 2001; Erlandson, et al. 2019; Merzendorfer and Zimoch 2003; Zhu, et al. 2016). Genes related to chitin metabolism are also among the fastest evolving functional categories in bumblebees, both in copy number variation and in sequence divergence (Figure 3; Additional file 2: Figure S15; Additional file 1: Table S18-19). These variable patterns of chitin-related gene evolution potentially underlie observed differences in morphology and insecticide resistance, which could influence the suitability of different species for commercial use. Across bumblebee genomes the fastest evolving genes are also related to processes including protein glycosylation, methylation, proteolysis, and tRNA aminoacylation for protein translation (Figure 3; Additional file 2: Figure S15; Additional file 1: Table S18-19). Protein glycosylation is involved in multiple physiological processes including growth, development, circadian rhythms, immunity, and fertility (Walski, et al. 2017). tRNA aminoacylation for protein translation process are involved in response to the changing environment (Pan 2013). Some genes that are not among the fastest evolving categories—for example, immune and detoxification genes, which are involved in the interaction of bumblebees with external environments—show differential patterns of positive selection in subsets of species (Additional file 1: Table S28; Additional file 1: Table S30), which can lead to gene functional divergence. Taken together, identification of the fastest evolving genes and those showing patterns of differential positive selection reveals substantial genetic variation across bumblebees. Future experimental investigations will be required to determine how the identified genetic variation is linked to specific differences in traits such as food preference, morphogenesis, insecticide and pathogen resistance, and the response to changing environments.

In addition to our discoveries regarding protein-coding genes, we found that TE-related sequences likely contribute to the variation of coding and regulatory repertoires (Figure 1; Additional file 1: Table S10-12). Compared with *non-Mendacibombus* bumblebees, *Mendacibombus* species have smaller genomes (Figure 1) and relatively narrow geographical distributions (Williams, et al. 2016). Considering TEs are the major determinant of genome size difference, with evidence that they were domesticated in bumblebee genomes, TEs may be implicated in the dispersal of non-*Mendacibombus* species across the globe, as they have been in other taxa (Baduel, et al. 2019; Casacuberta and González 2013; Schrader and Schmitz 2019).

More recent range expansions or contractions are driven, at least in part, by global climate change. To survive, bumblebees may have to move northward or to higher elevations as the climate warms (Kerr, et al. 2015; Soroye, et al. 2020). The sequenced genomes of species collected at high-elevation sites (> 4000 m) and others collected at low elevations (< 2000 m) (Figure 1) represent high quality genomic resources for investigating genes involved in high-elevation adaptation. We identified genes showing signs of positive selection in at least two subgenera of high-elevation species but not in any of the low-elevation species (Additional file 1: Table S31). These include genes putatively involved in eye development, muscle integrity maintenance, and metabolism, highlighting the importance of successful food-searching in high-elevation habitats where food is scarce. Exploring these further and identifying additional genomic features linked to life at high altitudes will help to understand differential successes of bumblebee species in a changing world.

## Conclusions

We have produced highly complete and accurate genome assemblies of 17 bumblebee species, including representatives from all of the 15 subgenera of *Bombus*. Our genus-wide comparative analysis of bumblebee genomes revealed how genome structures, genome contents, and gene evolutionary dynamics vary across bumblebees, and identified genetic variations that may underlie species trait differences in foraging, diet and metabolism, morphology and insecticide resistance, immunity and detoxification, as well as adaptations for life at high altitudes. Our work provides genomic resources that capture genetic and phenotypic variation, which should advance our understanding of bumblebee success and help identify potential threats. These resources form a foundation for future research, including resequencing and population genomics studies for functional gene positioning and cloning, which will inform the use of bumblebees in agriculture, as well as the design of strategies to prevent the decline of this important group of pollinators.

## Materials and Methods

### Sample collection and DNA extraction

Criteria including phylogenetic position, biological trait, geographic distribution, and specimen availability were applied to select species for whole genome sequencing. A total of 17 bumblebee species were selected (Additional file 1: Table S1), which span all of the 15 subgenera in the simplified classification system for the genus *Bombus* (Williams, et al. 2008). Among these, two species (*B. superbus* and *B. waltoni*) are from the subgenus *Mendacibombus*, which is sister to all other *Bombus* lineages; four species (*B. superbus*, *B. waltoni*, *B. skorikovi* and *B. difficillimus*) were collected at extremely high elevations (> 4000 m above sea level); two species (*B. turneri* and *B. skorikovi*) exhibit social parasitism; and one species (*B. polaris*) is endemic to the Arctic. In addition, species traits (i.e. range size, tongue length, parasite incidence, and decline status) vary across the selected bumblebees (Arbetman, et al. 2017). Samples were collected in the summer of 2016, with location and elevation information summarized in Additional file 1: Table S1. Their identities were confirmed by DNA barcoding as described (Hebert, et al. 2004). Genomic DNA was extracted from each specimen using the Gentra Puregene Tissue Kit (Qiagen). The abdomens of each sample were removed before DNA extraction to avoid microbial contamination.

### Genome sequencing and assembly

Genomic DNA purified from one single haploid drone of each species was used to generate one “fragment” library with an insert size of 400 or 450 bp using the NEBNext^®^ Ultra™ DNA Library Prep Kit for Illumina^®^ (NEB, USA). The prepared fragment libraries were sequenced on an Illumina HiSeq 2500 platform with a read length of 250 bp to produce overlapping paired-end shotgun reads (2 × 250 bp), and the target sequencing coverage was 100-fold or more for each species. Genomic DNA purified from multiple specimens of each species was used to generate four “jump” libraries (insert sizes: 4 kb, 6 kb, 8 kb, and 10 kb) according to reported methods (Heavens, et al. 2015). The prepared jump libraries were sequenced on an Illumina HiSeq X Ten platform, and paired-end reads (2 × 150 bp) were generated, with a sequencing depth of at least 40-fold coverage for each jump library. The sequencing results of “fragment” and “jump” libraries are summarized in Additional file 1: Table S2.

For each species, the 250 bp overlapping paired-end shotgun reads from the fragment library were processed using the software Seqtk (https://github.com/lh3/seqtk) to randomly subsample read pairs to achieve the total sequence length equivalent to ~60-fold sequencing coverage, a coverage recommended by the assembler we used (https://software.broadinstitute.org/software/discovar/blog/). Then, the subsampled shotgun reads were assembled using the software DISCOVAR *de novo* (version 52488), which performs well in assembling insect genomes (Love, et al. 2016), to produce contiguous sequences (contigs) for each species. Finally, shotgun reads from jump libraries were used to scaffold the contigs using the software BESST (Version 2.2.6) (Sahlin, et al. 2014). The obtained genome assemblies were checked for DNA contamination by searching against the NCBI non-redundant nucleotide database (Nt) using BLASTN (Camacho, et al. 2009), with an E-value cutoff of 1e-5.

To evaluate the quality and completeness of the genome assemblies, we compared genes present in the assemblies to a set of 4,415 universal single-copy orthologs (lineage dataset: hymenoptera_odb9) using the software BUSCO v3 (Waterhouse, et al. 2018).

### Genome annotation

#### RNA extraction and sequencing

For each species (*B. superbus, B. waltoni, B. confusus, B. soroeensis, B. consobrinus, B. difficillimus, B. haemorrhoidalis, B. turneri, B. opulentus, B. picipes, B. ignitus, B. sibiricus, B. breviceps,* and *B. pyrosoma*), total RNA was isolated using the TRIzol reagent (Invitrogen, CA, USA) following the manufacturer’s instructions. RNA integrity was evaluated on a 1.0 % agarose gel stained with ethidium bromide. After quantifying the concentration of RNA using a Qubit^®^ 2.0 Fluorometer (Life Technologies, CA, USA), 3 μg of RNA from each species was used to prepare sequencing libraries using the NEBNext^®^ UltraTM RNA Library Prep Kit for Illumina^®^ (NEB, USA) following manufacturer’s instructions. Library quality was assessed on the Agilent Bioanalyzer 2100 system. The prepared libraries were sequenced on the Illumina HiSeq X Ten platform, generating paired-end reads with a read length of 150 bp.

#### Protein-coding gene annotation

Annotation of protein-coding genes was based on *ab initio* gene predictions, transcript evidence, and homologous protein evidence, all of which were implemented in the MAKER computational pipeline (Cantarel, et al. 2008). Briefly, RNA-seq samples were assembled using Trinity (Haas, et al. 2013) with two different strategies using default parameters, *de novo* assembly and genome-guided assembly. Assembled transcripts were inspected by calculation of FPKM (fragments per kilobase of exon per million fragments mapped) expression values and removed if FPKM <1 and iso-percentage <3%. The filtered transcripts were imported into the PASA program (Haas, et al. 2003) for construction of comprehensive transcripts, as PASA is able to take advantage of the high sensitivity of reference-based assembly while leveraging the ability of *de novo* annotation to detect novel transcripts. The nearly “full-length” transcripts selected from PASA-assembled transcripts were imported to data training programs including SNAP (Korf 2004), GENEMARK (Lomsadze, et al. 2005) and AUGUSTUS (Stanke, et al. 2006). Afterwards, the MAKER pipeline was used to integrate multiple tiers of coding evidence and generate a comprehensive set of protein-coding genes.

The second round of MAKER was run to improve gene annotation. The predicted gene models with AED scores less than 0.2 were extracted for re-training using SNAP, GENEMARK, and AUGUSTUS. In addition, the RNA-seq reads were mapped to genomes using HiSAT2 and re-assembled using StringTie (Pertea, et al. 2016). The assembled RNA-seq transcripts, along with proteins from bees (superfamily Apoidea) that are available in NCBI GenBank (last accessed on 01/28/2018), were imported into the MAKER pipeline to generate gene models, followed by manual curation of key gene families.

### Functional annotation of the obtained gene models

To obtain functional clues for the predicted gene models, protein sequences encoded by them were searched against the Uniprot-Swiss-Prot protein databases (last accessed on 01/28/2018) using the BLASTp algorithm implemented in BLAST suite v2.28 (Altschul, et al. 1990). In addition, protein domains and GO terms associated with gene models were identified by InterproScan-5 (Jones, et al. 2014).

To evaluate the quality and completeness of gene annotation, we compared protein sequences predicted from the genome assemblies to a set of 4,415 universal singlecopy orthologs (lineage dataset: hymenoptera_odb9) using the software BUSCO v3 (Waterhouse, et al. 2018).

### miRNA annotation

Hairpin sequences downloaded from miRBase (http://www.mirbase.org/) were aligned to each reference genome using BLASTN (Altschul, et al. 1990) with an evalue cut-off of 10-6. Results were further filtered based on alignment length (≥50nt) and sequence similarity (≥80%). Mature sequences from miRBase were then mapped against this set of selected BLASTN hits, using Patman (Prufer, et al. 2008) with parameters -g 0 -e 1 (no gaps, up to one mismatch). Only genomic hits where at least one mature microRNA could be mapped with these criteria were retained. These were treated as a set of putative homologous microRNA genes.

Small RNA reads of *B. terrestris* were mapped to these predicted homologous loci, with no gaps or mismatches allowed. Genomic loci with at least 10 mapped reads were then selected, showing coverage at both the 5’ and 3’ ends. The final set of high confidence microRNAs was obtained by selecting all loci with the expected hairpin secondary structure, as predicted by RNAfold from the ViennaRNA package (Hofacker 2009), as well as strong evidence of Drosha-Dicer processing from the (manually inspected) patterns of small-RNA read alignments.

### tRNA annotation

All of the bumblebee genomes were screened with tRNAScan-SE (Lowe and Eddy 1997) to identify tRNA genes, with default parameters.

The prediction of lncRNAs: protein-coding potential for RNA transcripts was predicted using two algorithms, LGC version 1.0 (Wang, et al. 2019) and CPAT version 2.0.0 (Wang, et al. 2013). LGC could be used in a cross-species manner and the algorithm was applied directly to bumblebees, while CPAT requires high-quality training data to build a species-specific model. Considering bumblebees do not have enough high-quality “coding” and “non-coding” transcripts to build a model, the prebuilt fly model in CPAT was used. All the predictions were performed on a Linux platform. RNA transcripts were deemed to be non-coding if they were consistently predicted to be non-coding by both LGC and CPAT.

### Gene synteny analysis

MCScanX (Wang, et al. 2012) was used to identify syntenic blocks, defined as regions with more than five collinear genes, between *B. terrestris*, a previously published bumblebee genome (Sadd, et al. 2015), and each of the newly sequenced bumblebees with default parameters to infer synteny contiguity.

### *De novo* identification and annotation of transposable elements (TEs) Methods based on TE structure

LTR retrotransposons of the bumblebee genomes were *de novo* identified and annotated by LTRharvest and LTRdigest (Ellinghaus, et al. 2008; Steinbiss, et al. 2009). The identified LTR retrotransposons were further classified with the PASTEC module of the REPET package (Hoede, et al. 2014). When identifying LTR retrotransposons, TSD length was set to 4-6 bp and the minimum similarity of LTRs was set to 85%; the four-nucleotide termini of each LTR retrotransposon was set as TG…,CA. LTR length was set to 100-6000 bp. For the post-processing of LTRdigest, pptlength was set to 10-30 bp, pbsoffset to 0-5 bp, and trans to Dm-tRNAs.fa. pHMMs were used to define protein domains taken from the Pfam database. Non-LTR retrotransposons of the bumblebee genomes were identified and characterized using MGEScan-non-LTR, with default parameters (Rho and Tang 2009).

DNA transposons were identified by TBLASTN of known DNA transposase sequences that are available in Repbase (https://www.girinst.org/repbase/) against the bumblebee genome sequences. All regions that produced significant hits (E-values <1E-10) were excised with 3 kb of flanking regions. The terminal inverted repeats of a DNA transposon were identified through a self-alignment of the excised sequence using NCBI-BLAST 2.

### Methods based on the repetitive nature of TEs

RepeatScout (Price, et al. 2005) was used to *de novo* identify repetitive sequences from bumblebee genomes, with default parameters. The obtained consensus sequences were classified by the PASTEC module of the REPET package (Hoede, et al. 2014). All of the repetitive sequences were classified into Class I (retrotransposons), Class II (DNA transposons), Potential Host Genes, SSR (Simple sequence repeats) and “noCat” (which means no classification was found).

### TE landscapes in the bumblebee genomes

First, CD-HIT-EST (version 4.6.6) (Li and Godzik 2006) was used to parse TE sequences that were *de novo* identified based on structure and repetitive nature with a sequence identity threshold of 0.9 (other parameters as default) to reduce TE redundancy for each bumblebee species. Then, the remaining TE sequences from all the bumblebee species were combined to produce a comprehensive TE library. Using this repeat library, each bumblebee genome was analyzed with RepeatMasker (http://www.repeatmasker.org) to yield a comprehensive summary of the TE landscape in each species using Cross_Match as the search engine (other parameters as default). The annotation files produced by RepeatMasker were processed by inhouse scripts to eliminate redundancy. Refined annotation files were used to determine the TE diversity and abundance within each species. Tandem repeats in each genome were identified by Tandem Repeat Finder (Benson 1999), implemented in RepeatMasker.

### TEs proliferated after the divergence of *Mendacibombus* from the other subgenera

The subgenus *Mendacibombus* forms the sister group to all of the other extant bumblebees, diverging near the Eocene-Oligocene boundary approximately 34 million years ago (Cameron, et al. 2007; Hines 2008; Williams and Paul 1985). If a TE is present in one non-*Mendacibombus* species, but is absent at the orthologous positions in both *Mendacibombus* species (*B. superbus* and *B. waltoni*), then the TE is inferred to have transposed sometime after the divergence of the species from *Mendacibombus*. To identify such TEs in each of the non-*Mendacibombus* species, first, pairwise whole-genome alignments between the target species and *B. superbus* were performed using the software LASTZ (Harris 2007). Then, based on the whole genome alignment results, TE insertion scanner (https://github.com/Adamtaranto/TE-insertion-scanner) was used to identify “alignment gaps” showing signatures of TE insertions in the genome of the target species, with “--maxInsert 50000 --minIdent 85 --minInsert 80” choices (other parameters set as default). Secondly, 200 bp of sequence flanking the identified TE-like insertion on either side were extracted from the genomic sequences of the target bumblebee species and combined into one sequence of 400 bp. Then, the flanking sequences were used as queries in BLASTn searches against the genomic sequence of *B. waltoni*, with an e-value cutoff of 1e-10. Hits spanning both sides of the TE-like insertion with a minimal length of 350 bp were considered as empty sites in *B. waltoni* genome. Finally, TE-like sequences that have identifiable orthologous empty sites in both of the two *Mendacibombus* species were RepeatMasked by the comprehensive TE library of bumblebees to confirm their TE identity.

### The age distribution of TE families in bumblebees

The consensus sequence of each TE family was constructed using RepeatScout (Price, et al. 2005) for each of the 19 bumblebee species; this consensus represents the TE family’s master gene (i.e. ancestral sequence). The obtained consensus sequences were used to produce a species-specific TE library. Using these libraries, each genome was masked with RepeatMasker. Percent divergences from consensus sequences reported by RepeatMasker were converted to nucleotide distance measures using the Jukes-Cantor formula to correct for multiple hits. To increase accuracy, analyses were limited to TE elements ≥80% identical to their respective consensus sequences, with a minimum length of 80 bp. Results were pooled into bins of single unit distances and represent summaries of TE class proliferation history. Because TEs evolve neutrally following insertion, the age of individual TEs can be approximated by measuring the sequence divergence from the ancestral consensus sequence and by applying a neutral substitution rate of 3.6 × 10^-9^ for bumblebee (Liu, et al. 2017).

### The genomic distribution of TEs in bumblebees

The genomic coordinates of TEs in each species were compared with the coordinates of protein-coding genes in the same species to identify TEs that resided within or near predicted genes. Only when there were > 50 bp of overlap between a TE and predicted CDS was a TE considered to be overlapping with a coding region. In *B. terrestris*, the coordinates of TEs, excluding those found in coding regions, were also compared with the coordinates of open chromatin regions detected by ATAC-seq (Zhao, et al. 2019) to identify TEs that may serve as regulatory sequences. Orthologous groups containing genes whose coding regions have TE-derived sequences were extracted, along with their *dN/dS* values (see Molecular evolution analysis on gene functional categories section) to check their *dN/dS* ratios to determine if they are under selective constraint.

### Orthology delineation across *Apis* and *Bombus*

The locally installed OrthoDB pipeline (http://www.orthodb.org/software; Kriventseva et al., 2015) was employed to define orthologous groups for proteins coming from 19 bumblebees and 4 honeybees. In addition to the 17 newly sequenced bumblebees from this study, the following previously annotated gene sets were downloaded: *B. terrestris* (GenBank assembly: Bter_1.0), *B. impatiens* (GenBank assembly: BIMP_2.0), *Apis mellifera* (GenBank assembly: Amel_4.5), *Apis cerana* (GenBank assembly: ACSNU-2.0), *Apis florea* (GenBank assembly: Aflo_1.0), and *Apis dorsata* (GenBank assembly: *Apis dorsata* 1.3). Only the longest isoform of each gene was used in orthology delineation. The orthoMCL program (Li, et al. 2003) was applied to the same protein dataset to confirm the results of the OrthoDB pipeline on lineage- and species-specific genes, and only genes determined as lineage- or species-specific by both programs were used for downstream analysis. In order to characterize the function of *Bombus*-specific genes, genes from *B. terrestris* that are *Bombus*-specific were selected. The GO annotations of *Bombus*-specific genes were assigned by InterproScan-5 (Jones, et al. 2014) and visualized on the WEGO website (http://wego.genomics.org.cn/; gene level 4) (Ye, et al. 2006).

To construct the phylogeny for these 23 species (19 bumblebees and 4 honeybees), universal single-copy orthologs delineated by the OrthoDB pipeline were isolated, and 3,617 single-copy orthologs were identified. Protein sequences from each of those universal single-copy orthologs were aligned with the software MAFFT (Katoh, et al. 2002), followed by alignment trimming with BMGE (Criscuolo and Gribaldo 2010). Trimmed alignments were concatenated for each species, respectively, resulting in 23 long super-sequences. The super-alignment contained 2,008,306 amino acids with 222,460 distinct alignment patterns. IQTree version 2.0 (Minh, et al. 2020b) was used to construct a maximum likelihood concatenated tree with the ultrafast bootstrap method (Hoang, et al. 2018). The best-fitting amino acid substitution model for each partition was selected by automatically by IQTree’s internal implementation of ModelFinder (Kalyaanamoorthy, et al. 2017). A time calibrated, ultrametric tree was produced by using a non-parametric rate smoothing approach (Sanderson 2003) along with a fossil calibration range of 65 My to 125 My for the divergence of *Apis* and *Bombus* (Hines 2008). To assess phylogenetic discordance among loci, gene trees for each single-copy orthologous group were also reconstructed with IQTree (Additional file 1: Table S5)(Minh, et al. 2020b). Of the 3,617 gene trees, 3,530 could confidently be rooted by the outgroup genus *Apis* to count topologies (Additional file 1: Table S6). Rooting was performed using Newick Utilities (Junier and Zdobnov 2010). Gene and site concordance factors (CF) were then calculated for each node in the species tree as implemented in IQTree (Minh, et al. 2020a).

The quartet-based species tree reconstruction program ASTRAL (Zhang, et al. 2018), which can account for ILS, was also used for building the species phylogeny. The ggtree R package was used to visualize trees (Yu, et al. 2017).

### Estimate of ancestral genome sizes

The genome assemblies produced in this study were highly complete (Additional file 2: Figure S1), and genome assembly sizes do not correlate with assembly contiguity (p = 0.973; Additional file 2: Figure S23). Thus, smaller genome size estimates are unlikely to be artifacts of incomplete genome assembly, and quality control during assembly ensured that larger genomes were not due to extrinsic DNA contamination. Therefore, the genome assembly sizes should reflect true differences across bumblebees. Genome assembly sizes of the 19 sequenced bumblebees and four honeybees were obtained from the current study and published genome assemblies: *B. terrestris* (GenBank assembly: Bter_1.0), *B. impatiens* (GenBank assembly: BIMP_2.0), *Apis mellifera* (GenBank assembly: Amel_4.5), *Apis cerana* (GenBank assembly: ACSNU-2.0), *Apis florea* (GenBank assembly: Aflo_1.1), and *Apis dorsata* (GenBank assembly: *Apis dorsata* 1.3). Genome sizes were mapped onto the phylogenetic tree estimated in this study (Figure 1A), and ancestral genome sizes of bumblebees were estimated using parsimony ancestral state reconstruction in Mesquite 3.51 (http://www.mesquiteproject.org), with honeybee genome sizes serving as the outgroup.

### Hi-C library construction, sequencing, and assembly

For *B. turneri*, library preparation was performed by Annoroad Gene Technology (http://en.annoroad.com) and mainly followed a protocol described previously (Belton, et al. 2012). Briefly, thorax muscles of wild-caught males were cross-linked by 2% formaldehyde solution at room temperature for 20 mins, and 2.5 M glycine was added to quench the crosslinking reaction. After grinding with liquid nitrogen, homogenized tissues were resuspended in 25 ml of extraction buffer I (10 mM Tris-HCl [pH 8.0], 5 mM β-mercaptoethanol, 0.4 M sucrose, 10 mM MgCl2, 0.1 mM phenylmethylsulfonyl fluoride [PMSF], and 1x protease inhibitor [Roche]), then filtered through miracloth (Calbiochem). The filtrate was centrifuged at 3,500g at 4°C for 20 min. The pellet was resuspended in 1 ml of extraction II (10 mM Tris-HCl [pH 8], 0.25 M sucrose, 10 mM MgCl2, 1% Triton X-100, 5 mM β-mercaptoethanol, 0.1 mM PMSF, and 1x protease inhibitor) and then centrifuged at 18,400g and 4 °C for 10 min. The pellet was resuspended in 300 μl of extraction buffer III (10 mM Tris-HCl, [pH 8.0], 1.7 M sucrose, 0.15% Triton X-100, 2 mM MgCl2, 5 mM β-mercaptoethanol, 0.1 mM PMSF, and 1 x protease inhibitor) and loaded on top of an equal amount of extraction buffer III, then centrifuged at 18,400g for 10 min. The supernatant was discarded and the pellet was washed twice by resuspending it in 500 μl of ice-cold 1x CutSmart buffer, followed by centrifuging the sample for 5 min at 2,500g. The nuclei were washed by 0.5 ml of 1 x restriction enzyme buffer and transferred to a safe-lock tube. Next, the chromatin was solubilized with dilute SDS and incubated at 65 °C for 10 min. After quenching the SDS with Triton X-100, overnight digestion was applied with a four-cutter restriction enzyme (400 units of MboI) at 37 °C on a rocking platform. The flowing steps include marking the DNA ends with biotin-14-dCTP and performing blunt-end ligation of crosslinked fragments. The proximal chromatin DNA was re-ligated by ligation enzyme. The nuclear complexes were reverse-crosslinked by incubating with proteinase K at 65 °C. DNA was purified by phenol–chloroform extraction. Biotin-C was removed from non-ligated fragment ends using T4 DNA polymerase. Fragments were sheared to a size of 100–500 bp by sonication. The fragment ends were repaired by the mixture of T4 DNA polymerase, T4 polynucleotide kinase, and Klenow DNA polymerase. Biotin-labeled Hi-C samples were specifically enriched using streptavidin magnetic beads. A-tailing of the fragment ends were added by Klenow (exo-) and Illumina paired-end sequencing adapters were added by ligation mix. Finally, Hi-C sequencing libraries were amplified by PCR (12-14 cycles) and sequenced on the Illumina HiSeq X Ten platform, generating paired-end reads (2 × 150 bp). The Juicer tool (Durand, et al. 2016) was applied to map Hi-C reads against the contig sequences of *B. turneri* using the BWA algorithm (Heng, et al. 2010) with default parameters. Mapped reads with MAPQ quality scores ≥ 30 were chosen for the next analysis. Then, the 3D-DNA pipeline (Dudchenko, et al. 2017) was applied to assemble the scaffold sequences to the chromosome level.

For *B. ignitus*, *B. pyrosoma*, *B. breviceps*, and *B. haemorrhoidalis*, the *in situ* Digestion-ligation-only Hi-C protocol was employed to generate Hi-C reads as described (Lin, et al. 2018). In brief, for each species, brain tissue of wild-caught workers was ground into homogenate. Treated the samples and filtered the precipitated cells. Cells were double cross-linked with formaldehyde with EGS (Thermo) and 1% formaldehyde (Sigma). After that, the remaining formaldehyde was sequestered with glycine. The cross-linked cells were subsequently lysed in lysis buffer and incubated at 50 °C for 5min, placed on ice immediately. After incubation, the nuclei were digested by MseI (NEB, 100 units/μl). After restriction enzyme digestion, MseI biotin linkers were ligated to the digested chromatin respectively. Made the nuclei fragment-end phosphorylation. Next, added T4 DNA ligase (Thermo) to reaction complexes. Ligation was performed at 20 °C for 2h with rotation at 15 r.p.m. Then, purifying the proximity ligation DNA. The purified products were digested by MmeI at 37 °C for 1 h. The digested DNA sample was subjected to electrophoresis in native PAGE gels and the specific 80-bp DLO Hi-C DNA fragments were excised and purified. Next, Illumina sequencing adaptors were ligated to the 80-bp DLO Hi-C DNA fragments. After biotin incubation, the ligated DNA fragments were used as template and amplified by PCR (fewer than 13 cycle) to construct the Illumina sequencing libraries.

Hi-C sequencing libraries were sequenced on the Illumina HiSeq X Ten platform, generating 150 bp reads. The length of the DNA constructs in the DLO Hi-C library is between 78 and 82 bp. The length of a full linker is 40 bp, and the lengths of the target DNA sequences on each side of the linker are 19-21 bp. A Java program was used to exclude the linker parts from the reads and the target DNA fragments were used for downstream analysis. The Juicer tool (Durand, et al. 2016) was applied to map obtained target sequences against the scaffold sequences of each species using the BWA algorithm (Heng, et al. 2010), selecting the ALN parameter (other parameters as default). Mapped reads with MAPQ quality scores ≥ 30 were chosen for the next analysis. Then, the 3D-DNA pipeline (Dudchenko, et al. 2017) was applied to assemble the scaffold sequences to the chromosome level.

The coordinates of genes within scaffold sequences were converted into coordinates on chromosome sequences for those five species.

### Macrosynteny search and visualization

First, the longest CDS for each gene, along with their coordinates, were prepared for the bumblebee species with chromosome-level assemblies (*B. ignitus*, *B. pyrosoma*, *B. breviceps*, *B. haemorrhoidalis*, *B. terrestris* and *B. turneri*). Then, pairwise comparisons were performed between *B. turneri* and each of the other species using MCscan in the JCVI tool kit (https://github.com/tanghaibao/jcvi; last accessed Dec 25, 2019) (Wang, et al. 2012) to identify and visualize macrosynteny.

### Evaluation of chromosomal evolution rates

Orthologous genes and their coordinates on chromosomes were used as anchors to evaluate rates of chromosomal evolution. Two sets of orthologous genes for each pair of species were grouped together to form a standard input for the GRIMM-Synteny program v. 2.02 (Tesler 2002). The genome of *B. terrestris* was used as a reference for pairwise comparisons with other species genomes. Chromosomes of different species with similar sets of genes were named chromosomal elements. The GRIMM-Synteny program was run with default settings and the rearrangement distances (the number of conserved synteny blocks and inversions) were summarized.

### Global gene family evolution analysis

In order to identify rapidly evolving gene families within *Bombus,* protein sequences from the following species were used: *B. superbus, B. confusus, B. soroeensis, B. consobrinus, B. difficillimus, B. haemorrhoidalis, B. turneri, B. opulentus, B. picipes, B. ignitus, B. polaris, B. cullumanus, B. sibiricus, B. breviceps, and B. pyrosoma* (one species per subgenus was selected to avoid over-sampling in any subgenus). To ensure that each gene was counted only once, only the longest isoform of each gene in each species was used. An all-vs-all BLAST (Altschul, et al. 1997) search was then performed on these filtered sequences. The resulting e-values from the search were used as the main clustering criterion for the MCL program to group proteins into gene families (Enright and J. 2002). This resulted in 24,137 clusters. All clusters only present in a single species or not present at the root of the tree were then removed, resulting in 13,828 gene families. A time calibrated, ultrametric tree (Additional file 2: Figure S11) was built by taking the inferred *Bombus* phylogeny and using a non-parametric rate smoothing approach (Sanderson 2003) along with a fossil calibration range of 65 My to 125 My for the divergence of *Apis* and *Bombus* (Hines 2008).

With the gene family data and ultrametric phylogeny as input, gene gain and loss rates (λ) were estimated with CAFE v3.0 (Han, et al. 2013). This version of CAFE is able to estimate the amount of assembly and annotation error (ε) present in the input data using a distribution across the observed gene family counts and a pseudo likelihood search. CAFE is then able to correct for this error and obtain a more accurate estimate of λ. The resulting ε value was about 0.05, which implies that 5% of gene families have observed counts that are not equal to their true counts. After correcting for this error rate, λ = 0.0036. Using the estimated λ value, CAFE infers ancestral gene counts and calculates p-values across the tree for each family to assess the significance of any gene family changes along a given branch. Those branches with low p-values are inferred to be rapidly evolving. A Fisher’s exact test was performed on GO terms for genes in families that are rapidly evolving on any lineage vs. all other families, with a false discovery rate of 0.01.

### Protein domain variation across bumblebees

Predicted protein sequences were analyzed by InterproScan-5 (Jones, et al. 2014) to identify InterPro domains in each bumblebee species. InterPro domain annotations across the 19 bumblebee species were used to identify protein domains exhibiting the highest variation in gene counts across bumblebees. A crude measure that highlights such variation in copy-number was computed as the standard deviation divided by the mean of the bumblebee gene counts matching a particular InterPro domain. Results were filtered to focus on abundant domains, which have more than 200 genes in total and more than five genes in each bumblebee species.

### Molecular evolution analysis on gene functional categories

#### Orthology delineation across bumblebees

In addition to the 17 newly sequenced bumblebees from this study, we downloaded the two previously annotated gene sets for *B. terrestris* and *B. impatiens* from Ensembl (http://metazoa.ensembl.org/index.html). Only the longest isoform of each gene was used for downstream analysis. Protein sequences from the 19 bumblebees were used to delineate orthologous groups by locally installed OrthoDB software (OrthoDB_soft_2.4.4) (http://www.orthodb.org/software).

#### Assignment of functional categories to each orthologous group

GO term(s) and InterPro domain(s) associated with each gene of the orthologous group were identified by InterproScan-5 (Jones, et al. 2014). A GO term or InterPro domain was assigned to this orthologous group if more than 60% of the genes in it were assigned this GO term or InterPro domain by InterproScan-5.

#### Evolutionary rate (amino acid sequence divergence) estimation for each orthologous group

Evolutionary rates were computed for each orthologous group as the average of inter-species identities normalized to the average identity of all interspecies best reciprocal hits, computed from pairwise Smith-Waterman alignments of protein sequences. The ‘evolrate’ program of the OrthoDB_soft_2.4.4 package was used to obtain these rates.

#### *dN/dS* ratio estimation for each orthologous group

To avoid biases related to duplication among lineages and out-paralog genes, only universal single-copy orthologous groups (scOGs) were used to estimate *dN/dS* ratios. Protein sequences of scOGs were aligned by MAFFT (Katoh, et al. 2002) and then used to inform CDS alignments to generate DNA codon alignments with the codon-aware PAL2NAL program (Suyama, et al. 2006). Next, the aligned CDSs were trimmed by Gblocks (Talavera and Castresana 2007), with “-t c” and other parameters as default. After trimming, only orthologs consisting of aligned sequences from all species with a minimum of 150 bp and less than 20% Ns were retained for downstream analysis, which are available on-line (ftp://download.big.ac.cn/bumblebee/bumblebee-single-copy-orthologs.tar.gz). Then, based on trimmed alignments, Maximum Likelihood trees were constructed for each of the orthologous groups using RAxML-NG (Kozlov, et al. 2019). Finally, PAML (Yang 2007) was used to calculate the *dN/dS* ratio for each orthologous group using its respective phylogenetic tree (codeml model=0, NSsites=0, ncatG=1).

#### Enrichment analysis of the slowest and fastest evolving genes

Assignment of GO terms and InterPro domains was biased towards slower-evolving, well-conserved genes (Additional file 2: Figure S14), so the fastest evolving genes are less likely to be functionally annotated. Comparing the top enriched functional categories in the slowest and fastest subsets of genes could complement the GO and InterPro analyses described above. Orthologous groups with evolutionary rates and *dN/dS* ratios less than the 20th percentile or greater than the 80th percentile were selected to represent the slowest and fastest gene sets, respectively (Additional file 2: Figure S16). Enrichment tests on GO Biological Processes and Molecular Functions were performed using Bioconductor’s GOstats hypergeometric test (Falcon and Gentleman 2007) and with the topGO (http://www.bioconductor.org/packages/release/bioc/html/topGO.html) implementations of the classic Fisher and the weighted Fisher tests. The background gene sets in each case were genes from all 19 bumblebee genomes that were classified into any orthologous group and were annotated with Biological Process or Molecular Function GO-terms. The results were combined using a conservative strategy: terms must appear significant with a p-value <0.05 for all three enrichment tests, and there must be more than five genes in the test set. Complementary enrichment analyses using topGO’s implementation of the Kolmogorov-Smirnov (KS) were performed using evolutionary feature metrics: evolutionary rate (as above); universality (the proportion of species with genes in each orthologous group); and three copy-number metrics (average copy-number, copy-number variation, and proportion of species with duplicates). Only Biological Process terms associated with at least 10 orthologous groups were assessed. The KS test uses the score distributions directly without having to specify any top or bottom cut-off as described above for the classic tests with the 20th and 80th percentiles. Results are presented for terms showing significantly higher or significantly lower score distributions (Additional file 1: Table S14; Additional file 1: Table S19).

### Detection of positive selection signatures

1. Single-copy orthologous groups search: orthologous groups containing focal genes, along with their *dN/dS* values, were extracted from the **Molecular evolution analysis on gene functional categories** section. To avoid biases related to duplication among lineages and out-paralog genes, only universal single-copy orthologous groups were kept for downstream analysis.
2. Multiple sequence alignment and *de novo* gene tree construction: The multiple alignment and Maximum Likelihood tree of each ortholog were taken from the **Molecular evolution analysis on gene functional categories** section.
3. aBSREL analysis: For each ortholog, signatures of positive diversifying selection were searched using the aBSREL algorithm (https://www.datamonkey.org), with the respective multiple sequence alignment and Maximum Likelihood tree. Branches with test p-values < 0.05 were considered to be under selection.

### Intron evolution

Orthologous groups delineated across 19 bumblebees and one honeybee (*A. mellifera*) (deduced from the **Orthology delineation across *Apis* and *Bombus*** section) were examined to select a total of 8,672 with near-universal single-copy orthologue distributions: requiring no more than two species with no orthologues and no more than two species with multi-copy orthologues. These were further filtered to exclude groups with genes for which annotation features did not match the protein sequence and groups where the orthologues from five or more of the 20 species were singlecoding-exon genes (i.e. no introns), leaving 7,394 groups for the analysis. The protein sequences of the orthologues for each group were FASTA formatted with header information containing intron/exon data required for analysis with Malin (Csűros 2008). Protein sequences for each group were aligned with MAFFT v7.310 (Kazutaka and Standley 2013) using the ‘--auto’ option. The resulting alignments were then processed (two rounds of re-alignment) by the IntronAlignment tool from the Malin suite with option ‘-matrix blosum62 -rep 2’. The species tree and alignments were loaded into the Malin analysis tool and reliable intron sites were defined as having at least five non-gap amino acid positions in the alignment before and after the site and unambiguous characters in at least 18 of the 20 species. This resulted in a total of 45,804 sites for the analysis which was performed using the Bootstrap Posterior Probability (BPP) approach of Malin, using rate models computed from the default starting model with default optimization parameters and with one gain and one loss level.

### Stop codon readthrough analysis

#### Whole genome alignments

Before multiple whole genome alignments, repetitive regions of the 19 bumblebee and 4 honeybee (*Apis mellifera*, *Apis cerana*, *Apis florea*, and *Apis dorsata*) genome assemblies were first masked to reduce the total number of potential genomic anchors formed by the many matches that occur among regions of repetitive DNA. For whole genome alignments of the 23 bees, Cactus (Paten, et al. 2011), a reference-free whole genome aligner, was used. The phylogeny of 23-species estimated in this study (Figure 1A), with branch lengths reflecting neutral substitutions per site, was used as the guide tree.

#### Stop codon readthrough analysis

Annotation version GCF_000214255.1 for *B. terrestris,* obtained from NCBI, was used. The phylogeny is the 23-species maximum likelihood phylogeny estimated in this study. PhyloCSF (Lin, et al. 2011) was run on the region between the annotated stop codon (“first stop codon”) and the next inframe stop codon (“second stop codon”) referred to as the “second open reading frame (ORF)”, excluding both the first and second stop codons, of all annotated transcripts whose coding region ends in a stop codon, grouping together sets of transcripts having the same second ORF. For transcripts lacking an annotated 3’UTR, or for which the 3’UTR does not extend up to the second stop codon, the transcript was extened along the DNA strand without splicing. PhyloCSF was run using the default “mle” strategy and “bls” option, using the 12flies parameters but substituting the 23-bees tree. PhyloCSF computes a log-likelihood of an alignment under coding and non-coding models of evolution. The model assumes independence of codons given that the region is coding or non-coding. However, scores of neighboring codons are not independent. To correct for that, PhyloCSF-Ψ (Lin, et al. 2011) calculates a log-likelihood of length-dependent normal distributions trained on actual coding and non-coding regions of various lengths. Coefficients for PhyloCSF-Ψ were trained using coding regions at the ends of coding ORFs and non-coding regions at the starts of third ORFs, as described in (Jungreis, et al. 2016). The coefficients we obtained for *B. terrestris* were:

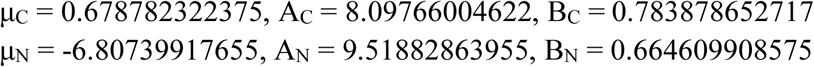

Both raw PhyloCSF scores and PhyloCSF-Ψ scores are reported in units of decibans. The 851 candidate readthrough stop codons in 817 genes were those satisfying all of the following conditions: (i) The second ORF is at least 10 codons long. (ii) PhyloCSF-Ψ > 0. (iii) The phylogenetic branch length of aligned species is more than 60% of the branch length of the full tree (enough to assure at least one *Apis* species is included). (iv) Species comprising at least 90% of the tree have the same first stop codon as *B. terrestris* (the *Drosophila* and *Anopheles* studies had found that readthrough stop codons are generally perfectly conserved). (v) Species comprising at least 60% of the tree have some stop codon aligned to the second stop codon. (vi) For second ORFs that overlap an annotated coding region on the same strand in the same reading frame, or on the opposite strand in the frame having the same third codon position (the “antisense” frame), the non-overlapping portion was required to be at least 10 codons long and have a positive PhyloCSF-Ψ score, as well as satisfying the branch length requirements described above.

To estimate the false discovery rate among our candidates, enrichment of the TGA stop codon with 3’ base C was used, which is known to be the “leakiest” 4-base stop codon context (Bonetti, et al. 1995) and is highly enriched among readthrough stop codons (Jungreis, et al. 2011). Of the 851 stop codons in the list, 172 (20.2%) have the TGA-C context, whereas of the 8059 annotated stop codons for which the second ORF has negative PhyloCSF-Ψ score and are thus unlikely to be readthrough, only 280 (3.5%) have the TGA-C context. Among the readthrough stop codons previously reported in *Drosophila* 32.2% had the TGA-C context (Jungreis, et al. 2011). If a similar fraction holds in *Bombus*, the number of actual readthrough stops codons among the 851 would be approximately (172 – 3.5% × 851) / (.322 –.035) = 496. Even if as many as 50% of readthrough stop codons in *Bombus* use TGA-C, a similar calculation provides a conservative estimate that the list includes 306 readthrough transcripts. Among the 200 of the candidates with highest PhyloCSF-Ψ score, 72 have TGA-C stop context, so a similar calculation conservatively estimates 140 readthrough transcripts among these 200 candidates, for a false discovery rate of no more than 30%.

### Codon usage bias analysis

Codon usage bias, the preferential use of specific synonymous codons, is a pattern maintained by mutation-selection-drift balance. The selection is linked to the efficiency and/or accuracy of translation. The selective effect of codon usage is only slightly advantageous and consequently selection’s efficiency depends on population size (Subramanian 2008; Vicario, et al. 2007); species with larger population sizes have more efficient selection for codon usage bias. Within the genomes, strength of selection could vary based on the stage of development when the genes are mainly translated (Vicario, et al. 2007). To determine the evolutionary forces affecting codon usage bias across bumblebees, a set of universal orthologous protein-coding genes was used (delineated in the **Molecular evolution analysis on gene functional categories** section). A total of 3,521 genes, which are present in all 19 species and have at least 50 unambiguous codons (no N or other ambiguity letters), were used for codon bias analysis. Candidate optimal codons were defined by examining the correlation between overall gene codon usage bias and the preference of use of a single codon as performed previously (Vicario, et al. 2007). As an estimator of overall codon usage bias, the Effective Number of Codons (ENC) was used, which was estimated by using the exponential of the sum of Shannon entropy of each codon family frequency set. As an estimator of preference for a single codon, the relative synonymous codon usage (RSCU) was used.

### Gene family evolution analysis of chemosensory genes

To detect the putative chemosensory genes of the three major gene families □ odorant receptors (ORs), gustatory receptors (GRs) and ionotropic receptors (IRs) □ from the 17 newly sequenced and the *B. impatiens* genomes, TBLASTN searches (with 1e-5 as the e-value cutoff) (Gertz, et al. 2006; Karpe, et al. 2016) were performed using the protein sequences of *A. mellifera* (Robertson and Wanner 2006) and *B. terrestris* (Sadd, et al. 2015) as queries. Putative chemosensory gene-containing regions were extracted from each genome to predict gene models using the protein2genome module of Exonerate v2.2.0 (Slater and Birney 2005). These putative gene-containing regions were separately re-examined if there were no good hits based on Exonerate. Candidate chemosensory genes were further manually refined and checked for the characteristic domains of ORs (IPR004117), GRs (IPR009318 or IPR013604), or IRs (IPR019594 or IPR001320) in their encoded protein sequences using InterProScan v5.27-66.0 (Jones, et al. 2014; Zhou, et al. 2015; Zhou, et al. 2012). Partial sequences were completed with the nearest START and/or STOP codons wherever possible. Probable amino acid sequences of pseudogenes, which were identified using in-frame STOP codons or frameshifts, were determined from their predicted coding regions, and the letter “X” was used to represent STOP codons and frameshifts. The letter “Z” denotes unknown amino acids. The same procedure was repeated, using newly identified chemosensory genes as queries, until no additional genes were found. Gene names were assigned following the closest homologue of *B. terrestris*. When there were two or more gene copies in one analyzed species but a single-copy in *B. terrestris*, candidate gene names were suffixed with a, b, c, and so on. For ORs and GRs, genes encoding intact proteins with a length >= 350 amino acids were kept for downstream analysis.

Multiple alignments of the available bumblebee chemosensory genes were generated using MAFFT v7.407 (“E-INS-i strategy) (Kazutaka and Standley 2013), poorly aligned regions in the alignments were filtered using TrimAl v1.4 (“automated1” option) (Capellagutierrez, et al. 2009), and maximum-likelihood phylogenetic trees were estimated using RAxML v8.2.11 (with the “PROTCATJTTF” model and 100 bootstrap replicates) (Stamatakis 2014). To estimate the numbers of gains and losses of chemosensory genes, we used maximum-likelihood-based and parsimony-based approaches, respectively; all genes of each chemoreceptor family were used as input for CAFE v4.2 (De Bie, et al. 2006) with default settings, and gene trees were reconciled with species tree using Notung v2.9.1 (Chen, et al. 2000).

Signatures of positive selection were searched for OR, GR and IR genes as described in **Detection of positive selection signatures** section.

### Evolution of genes involved in detoxification

Glutathione-S-transferases (GSTs), carboxyl/cholinesterases (CCEs), and cytochrome P450 monooxygenases (P450s) are involved in the detoxification of xenobiotics. To identify detoxication genes in the newly sequenced bumblebees, annotated P450, GST, and CCE protein sequences of *B. terrestris, A. mellifera,* and *D. melanogaster* were used as queries to search against the predicted protein sequences from each genome using BLASTp (Altschul, et al. 1990). If certain genes appeared to be missing, TBLASTn was used as in annotating chemosensory genes. All of the identified detoxication genes were further checked for the presence of their characteristic domains to confirm their identity (GST [IPR004045 and IPR010987], P450 [IPR001128], and CCE [IPR002018]).

Signatures of positive diversifying selection were searched for each category of detoxication genes as described in **Detection of positive selection signatures** section.

### Identification and characterization of immune genes

To identify immune-related genes in the newly sequenced bumblebees, annotated immune genes of *B. terrestris* and *A. mellifera* were used as queries to search against the predicted protein sequences from each genome using BLASTp (Altschul, et al. 1990). If certain genes appeared to be missing, TBLASTn was used as in annotating chemosensation genes.

Immune genes were classified into three broad functional categories — “recognition,” “signaling,” and “effector” — based on previous reports (Barribeau, et al. 2015; Evans, et al. 2006; Neafsey, et al. 2015; Sackton, et al. 2007; Waterhouse, et al. 2020). Specifically, the recognition class includes SCR (scavenger receptors), GNBP (gram-negative binding proteins), PGRP (peptidoglycan recognition proteins), and GALE (galectins). The signaling class includes TOLL (toll-like receptors), JAKSTAT (Jak/Stat pathway members), IMDPATH (Imd pathway members), CLIP (CLIP-domain serine proteases), SRPN (serine protease inhibitors), CASP (caspases), and IAP (inhibitors of apoptosis). The effector class includes SOD (superoxide dismutases), TEP (thioester-containing proteins), LYS (lysozymes), PPO (prophenoloxidases), PRDX (peroxidases), AMP (anti-microbial peptides), ML (MD2-like proteins), NIMROD (nimrod-related proteins), FREP (fibrinogen-related proteins), and CTL (C-type lectins).

Signatures of positive diversifying selection were searched for each category of immune genes as described in **Detection of positive selection signatures** section.

### Evolutionary analysis of sex-determination genes

Protein sequences of *B. terrestris* genes including *feminizer (fem), feminizer* 1 (*fem* 1), and *transformer 2,* which are involved in the sex determination pathway, were used as queries to search against the newly sequenced genomes by locally installed BLAST (Gertz, et al. 2006) to identify their orthologs/paralogs in bumblebees. Before phylogenetic analysis, sequences were multiply aligned using MUSCLE (Edgar 2004). The evolutionary history of sex-determining genes in *Bombus* and related species was inferred using Maximum Likelihood with the JTT matrix-based model implemented in MEGA X (Jones, et al. 1992; Kumar, et al. 2018). The tree with the highest log likelihood (−6161.36) is shown. A discrete Gamma distribution was used to model evolutionary rate differences among sites (5 categories (+G, parameter = 2.2)) with branch lengths measured in the number of amino acid substitutions per site. RELAX (Wertheim, et al. 2015) was employed to test whether the strength of natural selection was relaxed or intensified along a specified set of test branches. The spurious action of diversifying selection in a subset of branches was detected by aBSREL (Smith, et al. 2015). To further identify individual sites that were subject to episodic diversifying selection, the mixed effect model of evolution (MEME) was applied (Murrell, et al. 2012).

### Identification of genes involved in the adaptation of bumblebees to high elevation

To identify genes involved in high-elevation adaptation, searches were conducted for genes undergoing positive selection in *B. superbus*, *B. waltoni*, *B. difficillimus*, and *B. skorikovi*, which were all collected at elevations > 4,000 m (Figure 1). First, universal single-copy orthologous groups were obtained, along with their respective multiple sequence alignments and Maximum Likelihood trees (described in the **Molecular evolution analysis on gene functional categories** section). Then, the improved branch-site model in the Codeml program of the PAML package was used to identify genes showing signatures of positive selection (Zhang, et al. 2005). In brief, *B. superbus*, *B. waltoni*, *B. difficillimus*, and *B. skorikovi* (all collected at elevations > 4,000 m) were assigned as the foreground branches and all the other bumblebee species (all collected at elevations < 2,000 m) as the background branches. A positive selection model that allowed a class of codons on the foreground branches to have *dN/dS* > 1 (model = 2, NSsites = 2, omega = 0.5|1.5, fix_omega = 0) was compared with a null model that constrained this class of sites to have *dN/dS* = 1 (model = 2, NSsites = 2, omega = 1, fix_omega = 1) using a likelihood ratio test and calculated a p-value for each comparison. Multiple comparisons were corrected for by using the Benjamini and Hochberg method and selected genes with an adjusted p-value < 0.05 as candidate positively selected genes (PSGs). Then, the Bayes Empirical Bayes (BEB) method (Yang, et al. 2005) was used to calculate posterior probabilities for site classes to identify codon positions that experienced positive selections (*dN/dS* > 1). Candidate PSGs that also contained codon positions showing significant BEB values (posterior probability >95%) were further analyzed using the software aBSREL (Smith, et al. 2015) to identify genes that show positive selection in at least two subgenera of high-elevation species but not in any of the low-elevation species. Such genes were believed to be PSGs involved in high-elevation adaptation. Finally, Codeml was used to estimate *dN, dS,* and *dN/dS* of these PSGs with the free ratio model (model = 1, NSsites = 0). PSGs with *dS* >1, suggesting considerable saturation at the synonymous sites, were removed from downstream analysis to avoid false positives. Functional clues about the identified PSGs were obtained by BLAST searching against the UniProt database (https://www.uniprot.org) and by literature review.

### Evolution of piRNA genes

Protein sequences for *Ago1, Armitage, Eggless, Gasz, Hen1, Maelstrom, Minotaur, Papi*, *Piwi/Aub*, *Qin*, *Shutdown*, *Spindle-E*, *Squash*, and *Trimmer* in *Apis mellifera* were downloaded from GenBank based on the dataset used by (Wang, et al. 2017). A BLAST protein database was built from the transcriptomes of each *Bombus* species and selected the top BLASTp hits for each species. We restricted our analyses to proteins that were present and had a single copy for all of the species.

Protein sequences were aligned using PSY-Coffee and automatically trimmed using G-Blocks while allowing for smaller final blocks and gap positions within the final blocks (Notredame, et al. 2000; Talavera and Castresana 2007). Phylogenies were estimated in MrBayes 3.2 with *Apis mellifera* set as the outgroup (Ronquist, et al. 2012). A mixed model for amino acid evolution was used. Each analysis ran for 10 million generations with the sampling frequency set to 1,000 with 3 heated chains, and 25% of the trees discarded as burnin.

The trimmed multiple alignments of single-copy orthologous groups containing piRNA genes, along with their phylogenies, were extracted from **Molecular evolution analysis on gene functional categories** section. Positive selection was detected by aBSREL (Smith, et al. 2015).

However, analysis of branch lengths and positive selection for 14 piRNA pathway genes across bumblebees found neither to be associated with genome size.

