## Additional file 2 for "Genus-wide characterization of bumblebee genomes reveals variation associated with key ecological and behavioral traits of pollinators"

### Figure S1: BUSCO assessment of genome assembly completeness.

A set of 4,415 universal single-copy orthologs (lineage dataset: hymenoptera_odb9) were used to check their presence and completeness in each of the 17 genome assemblies.

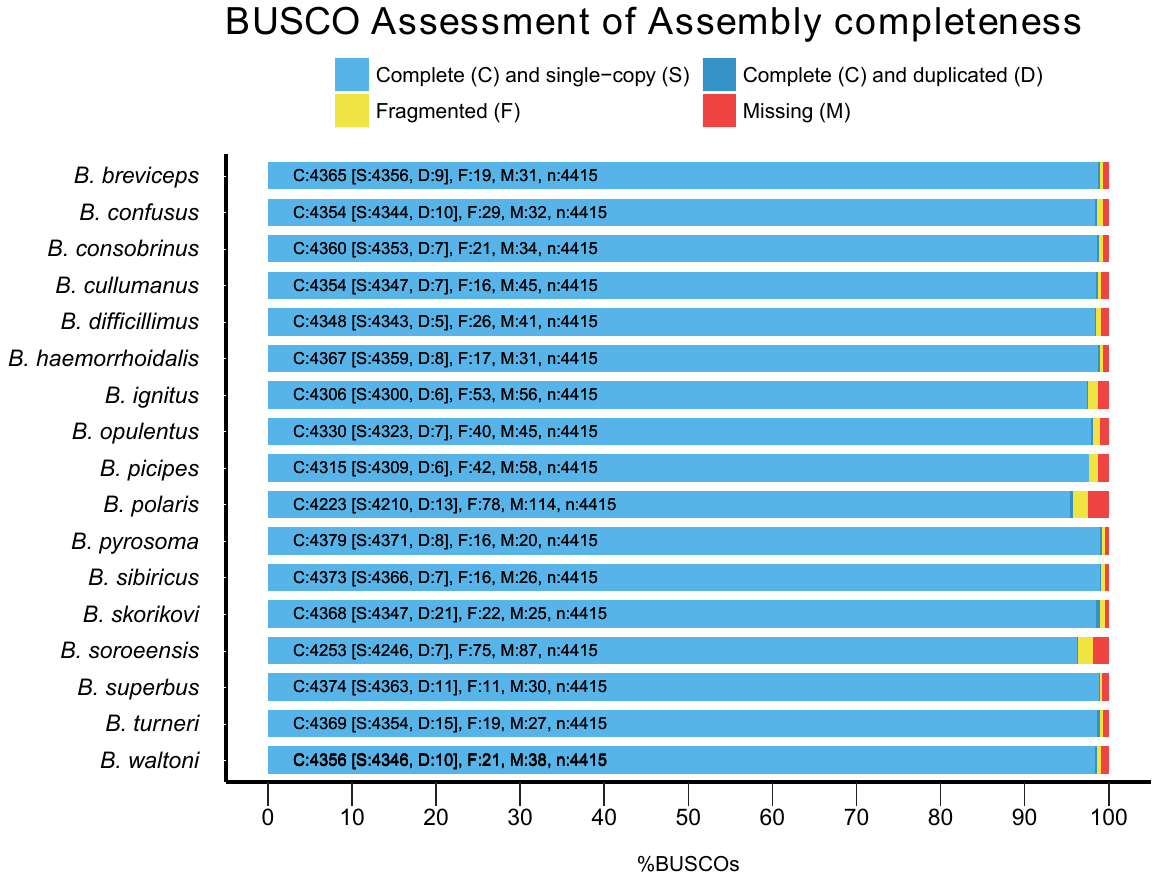

### Figure S2: Correlation between gene count and genome assembly contiguity.

Pearson correlation analysis between gene count and genome assembly contiguity (scaffold N50) of the 17 newly produced bumblebee genome assemblies.

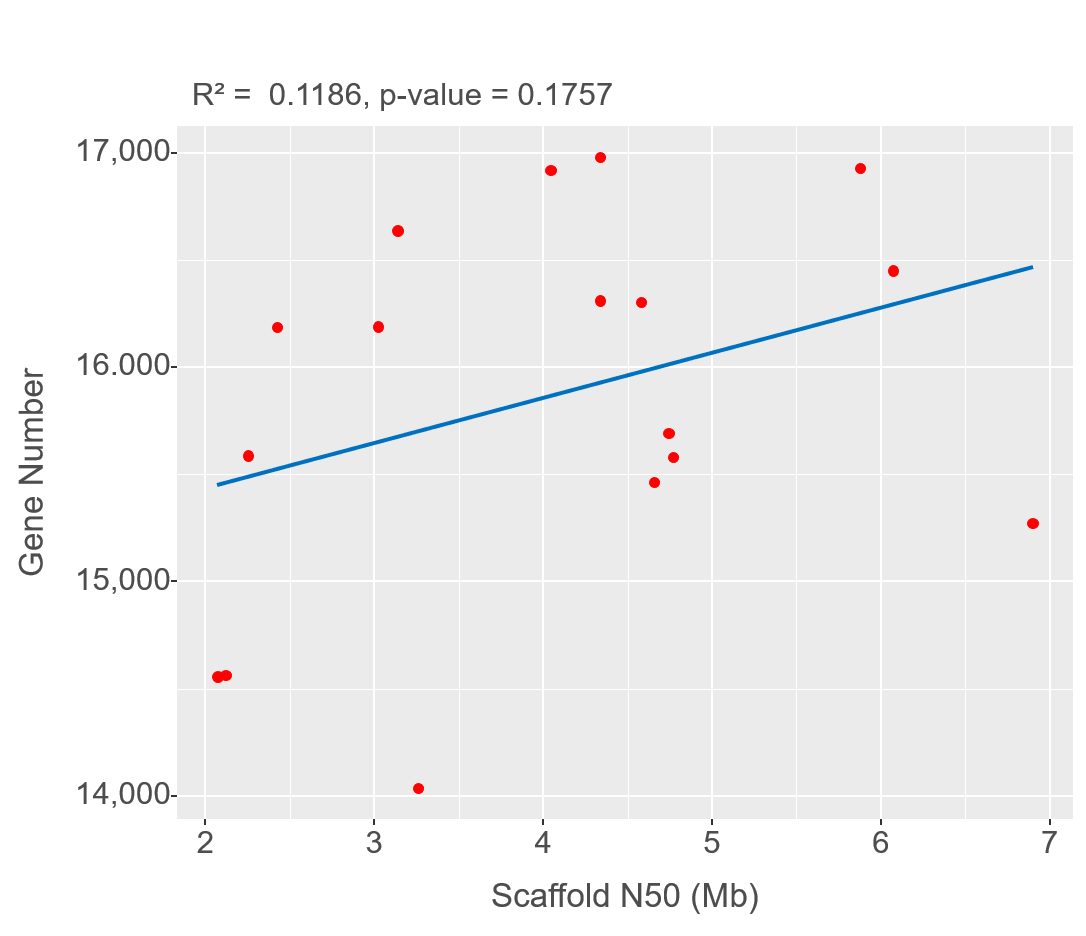

### Figure S3: BUSCO assessment of genome annotation quality.

A set of 4,415 universal single-copy orthologs (lineage dataset: hymenoptera_odb9) were used to check their presence and completeness in each of the 17 predicted proteomes.

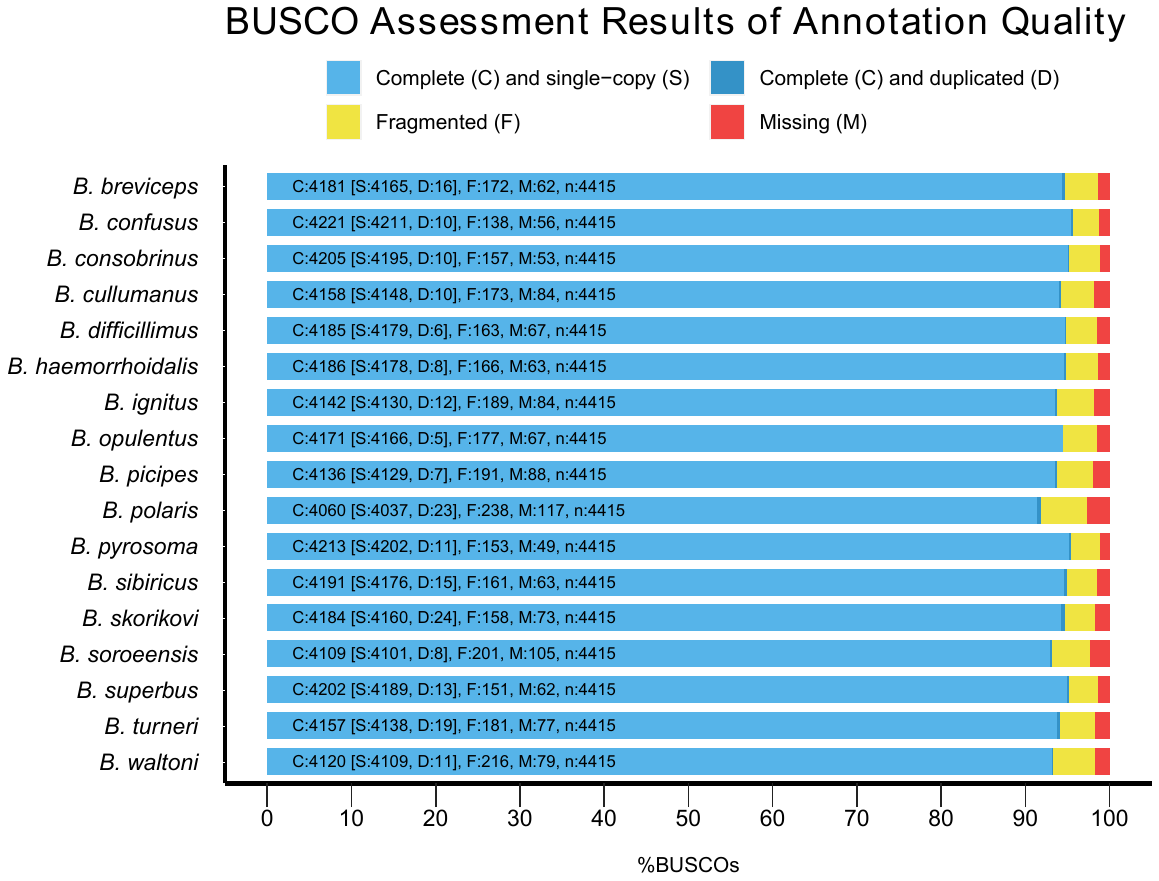

### Figure S4: Comparison of maximum likelihood concatenated and quartet-based (ASTRAL) species topologies.

Branch lengths are unscaled, with numbers beside each node representing bootstrap values. The only topological difference is highlighted in the teal rectangle.

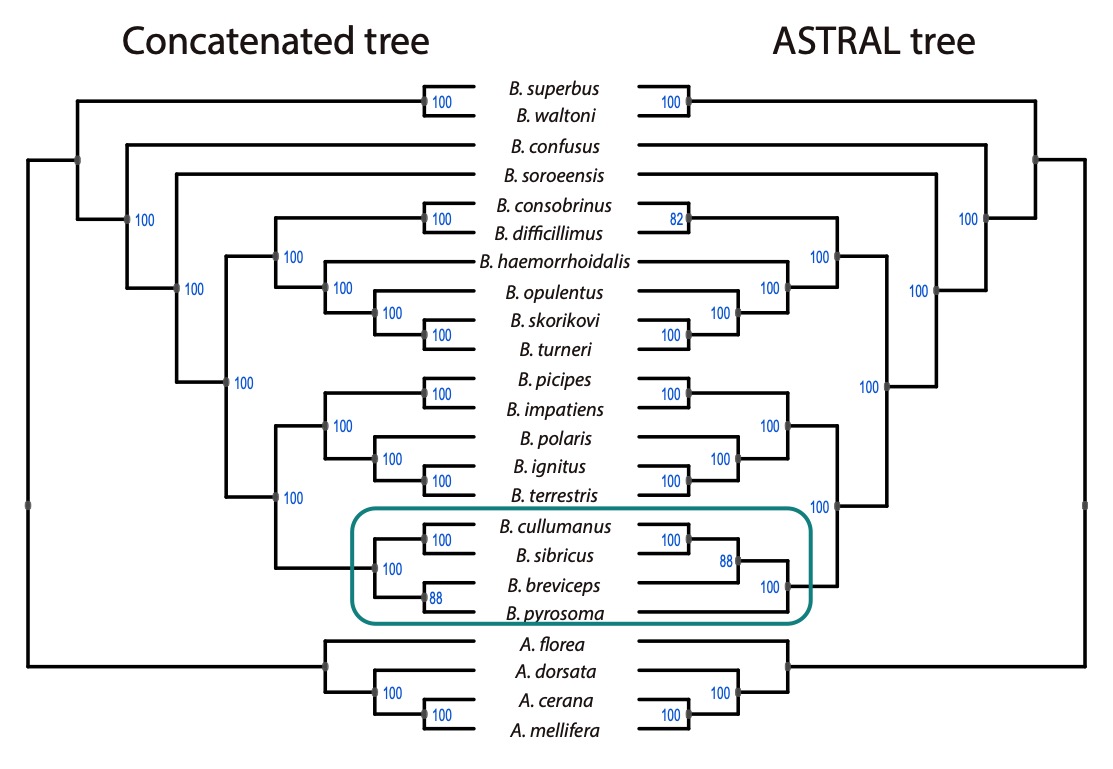

### Figure S5: Branch length and concordance factor are highly correlated in both the concatenated (A) and ASTRAL (B) trees.

The points each represent one internal node and the dashed line is the best-fit line of a linear regression. Branch lengths in the maximum likelihood concatenated tree represent relative numbers of substitutions while branch lengths in the ASTRAL tree represent coalescent units.

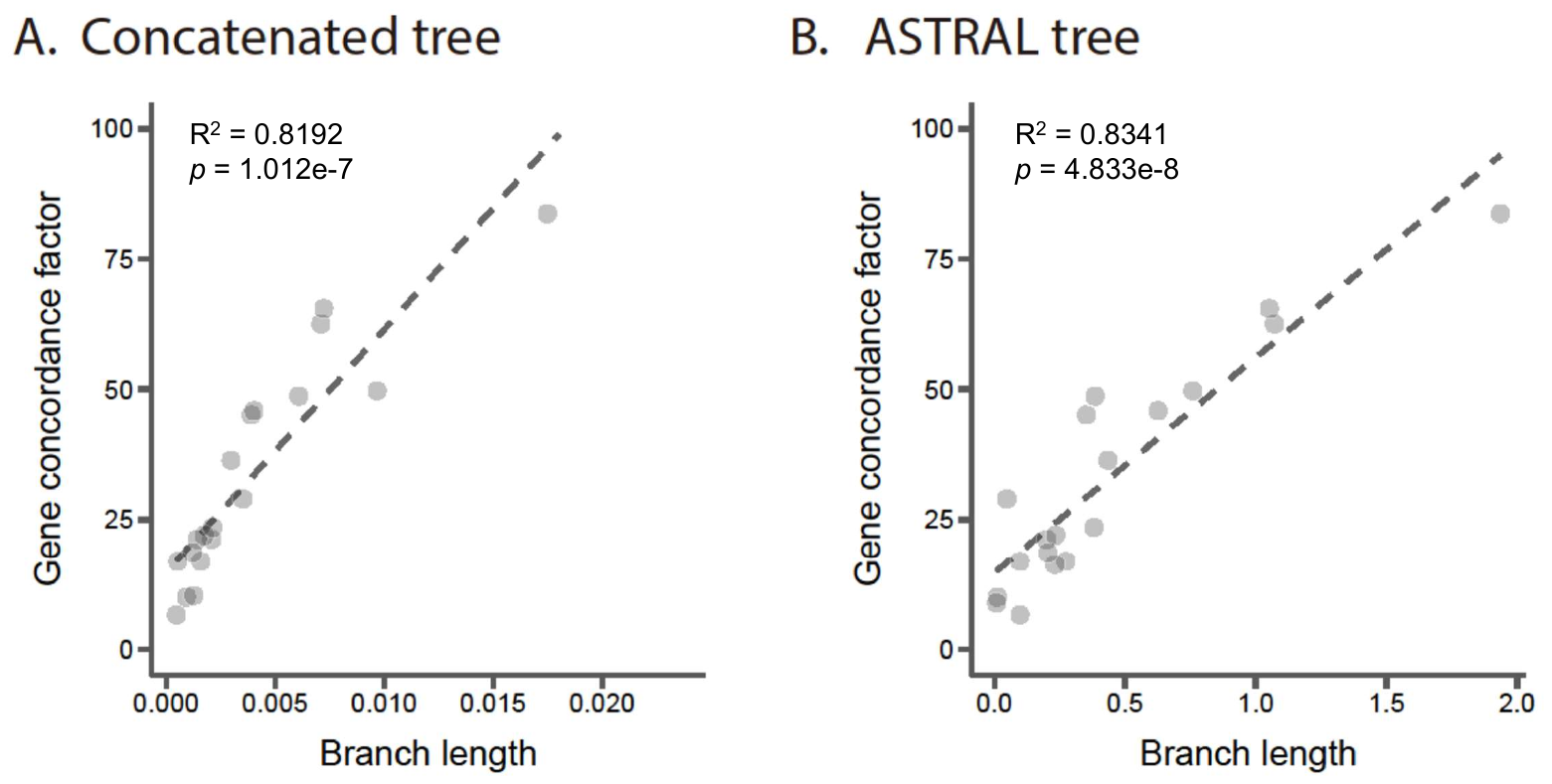

### Figure S6: Hi-C assisted genome assembly and the origin of *B. turneri* chromosomes.

Hi-C contact heatmaps for *B. breviceps* (**A**) and *B. ignitus* (**B**). (**C**) Macrosynteny across *B. breviceps*, *B. turneri* and *B. ignitus* to show the origin of *B. turneri* chromosomes. (**D**). Macrosynteny between *B. turneri* and subgenus *Bombus* species (*B. ignitus* and *B. terrestris*) to show the origin of *B. turneri* chromosomes.

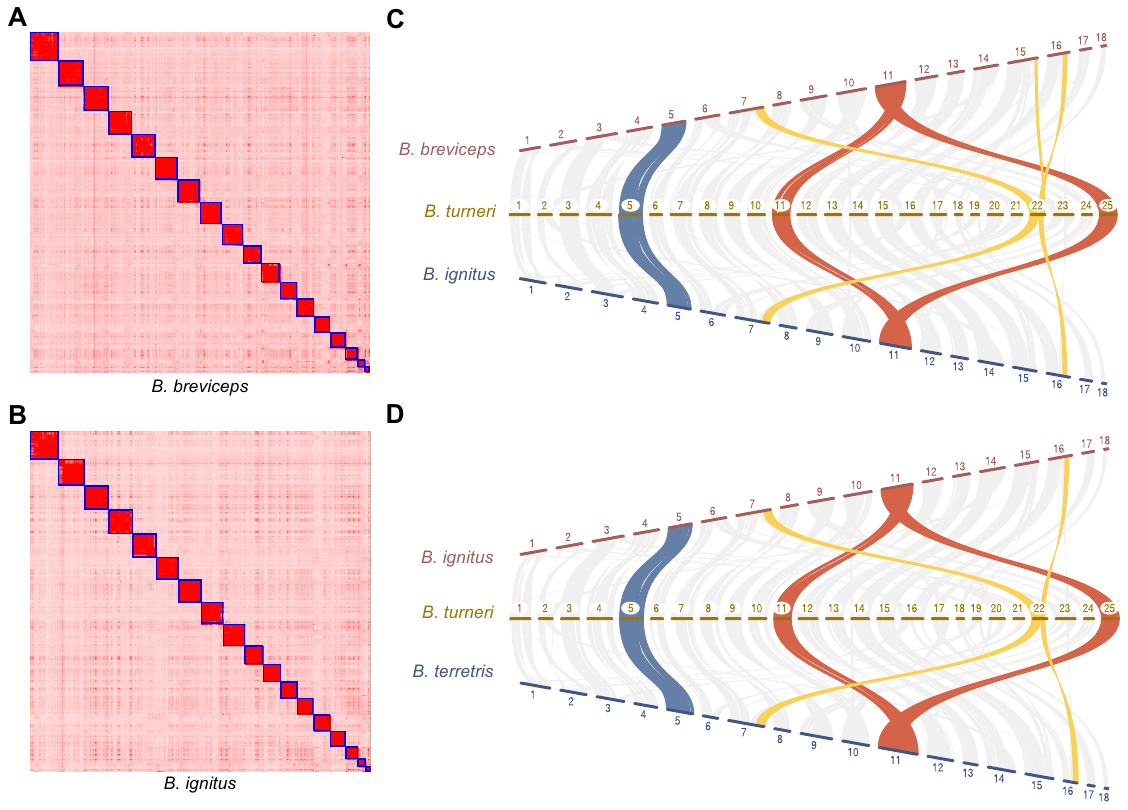

### Figure S7: Ancestral genome size of bumblebees inferred by Mesquite 3.51.

Numbers on nodes indicate the inferred ancestral genome sizes (in Mb). The genome assembly sizes of each species were shown in brackets following their species names (in Mb).

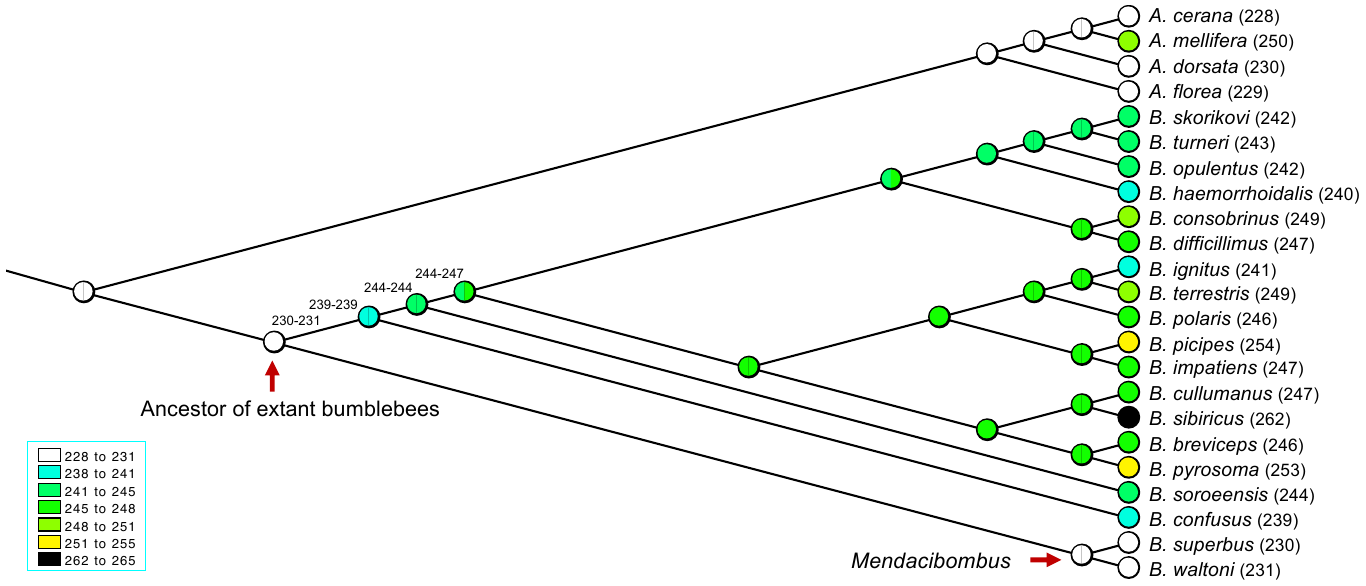

### Figure S8: Simple sequence repeat content versus genome size differences.

**Pearson correlation analysis between differences in simple sequence repeat content relative to that of *B. superbus* and differences in genome size (relative to that of *B. superbus*).**

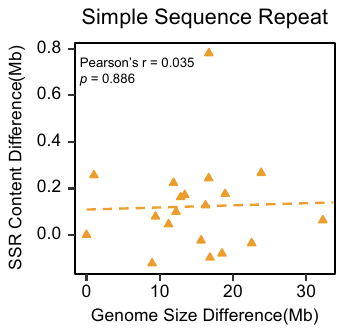

### Figure S9: Transposable element counts.

**The number of TEs in each non-*Mendacibombus* species that proliferated after the divergence of their host species from *Mendacibombus* species.**

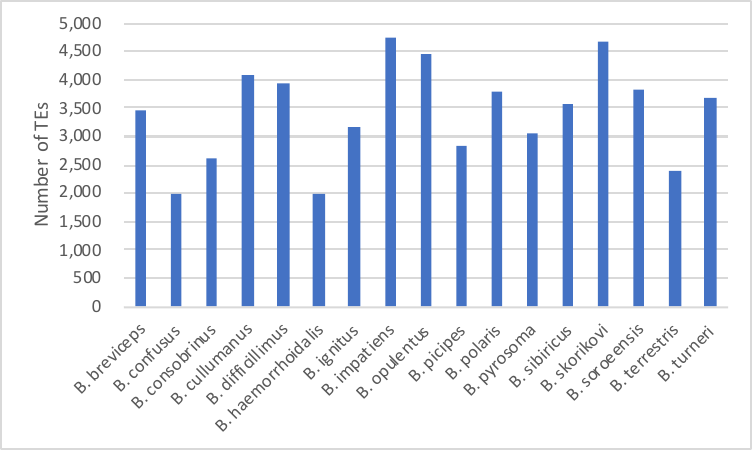

### Figure S10: TE proliferation history in *Mendacibombus* species (*B. superbus* and *B. waltoni*) and in two representative non-*Mendacibombus* species (*B. terrestris* and *B. turneri*).

Red arrows indicate the amplification peak of *Mendacibombus* species in sequence divergence from consensus sequences and in Million years ago (Mya), respectively. Non-*Mendacibombus* species (*B. terrestris* and *B. turneri*) have a more recent amplification peak than that of *Mendacibombus* species (*B. superbus* and *B. waltoni*).

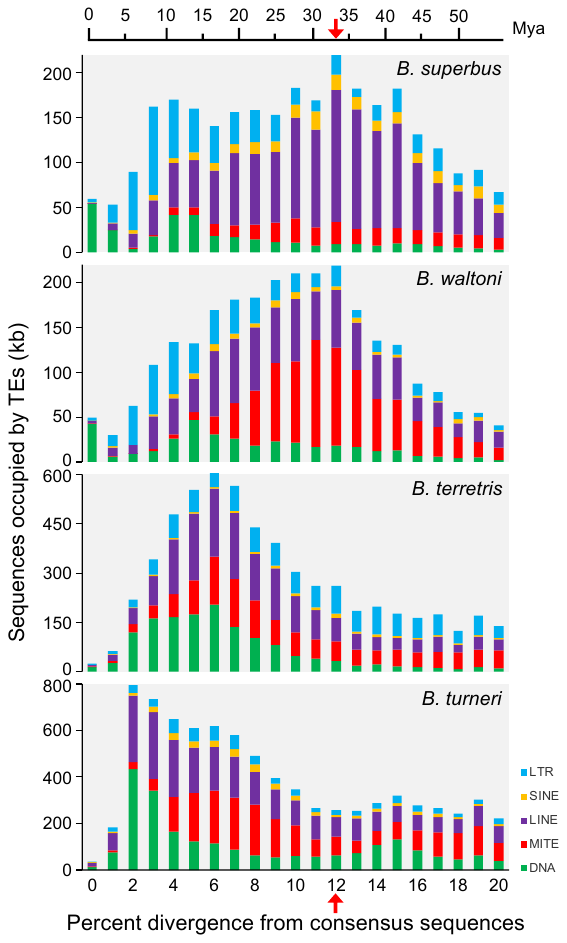

### Figure S11: Rates of gene gain/loss across the phylogeny of *Bombus*.

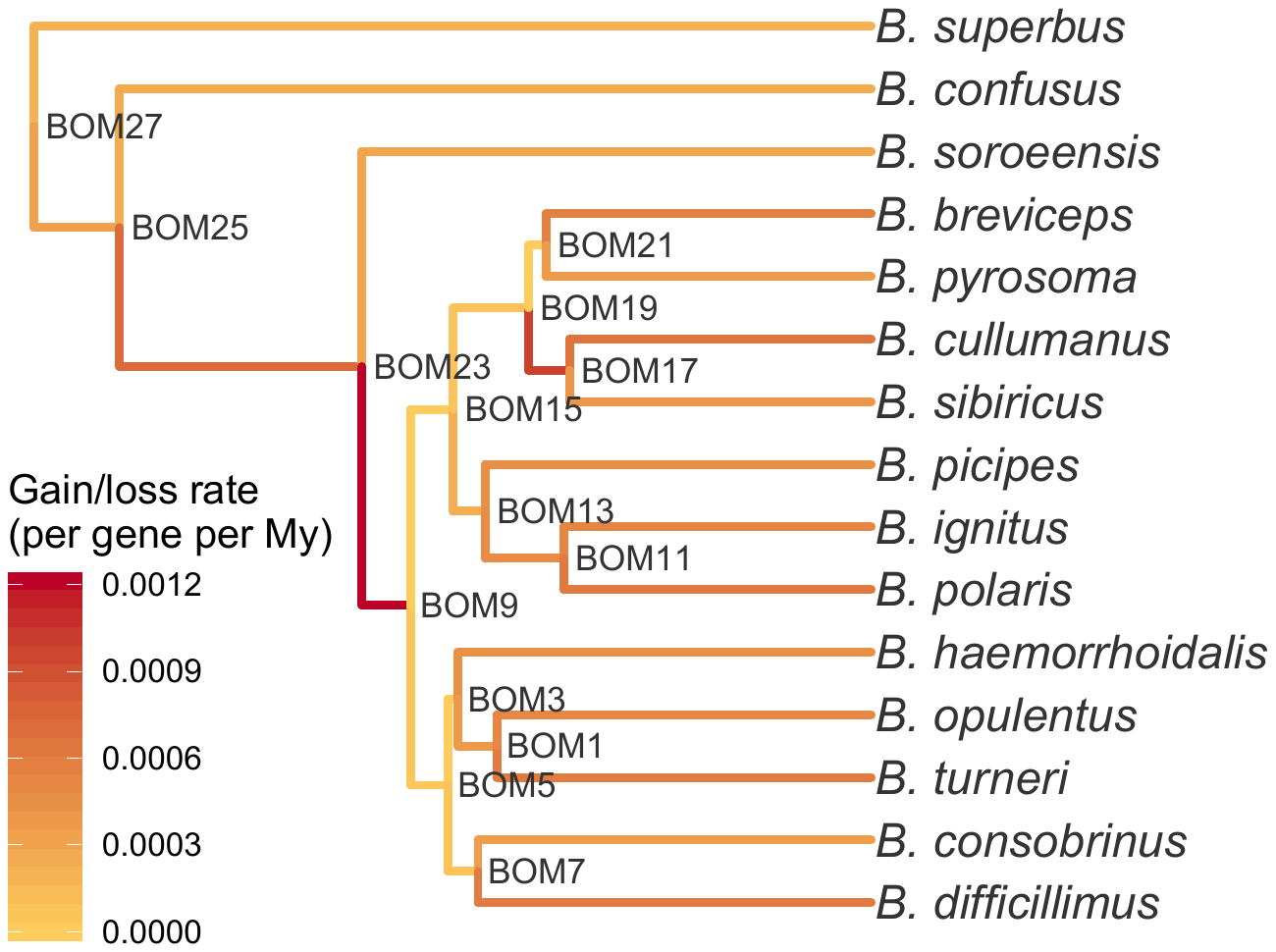

### Figure S12: One example of stop codon readthrough in *B. terrestris*.

Alignment of the readthrough region of transcript rna11916 in gene XM_012313001.2, color coded by CodAlignView (https://data.broadinstitute.org/compbio1/cav.php). Also shown are the third ORF, and 10 codons on each end. All substitutions in both the second and third ORFs are synonymous (light green), a strong indication that these regions are protein-coding, which would indicate that both the annotated TGA stop codon and the subsequence TAG stop codon are read through, making this a double-readthrough gene. After the TGA stop codon that ends the third ORF, there are many non-synonymous substitutions (red and dark green), frame shifting indels (grey and orange), and stop codons, typical of non-coding regions. Most protein-coding regions have some non-synonymous substitutions, so the readthrough extension of the rna11916 protein is unusually well conserved. The evolutionary coding potential as measured by PhyloCSF of the 38-codon second ORF (180.4) and the 57-codon third ORF (295.5) are more than 2000 times as likely to occur in coding regions than non-coding regions, implying that it these extensions have been functional at the amino acid level in much of the bee tree. The perfectly conserved TGA-C stop codon context is known to promote inefficient termination.

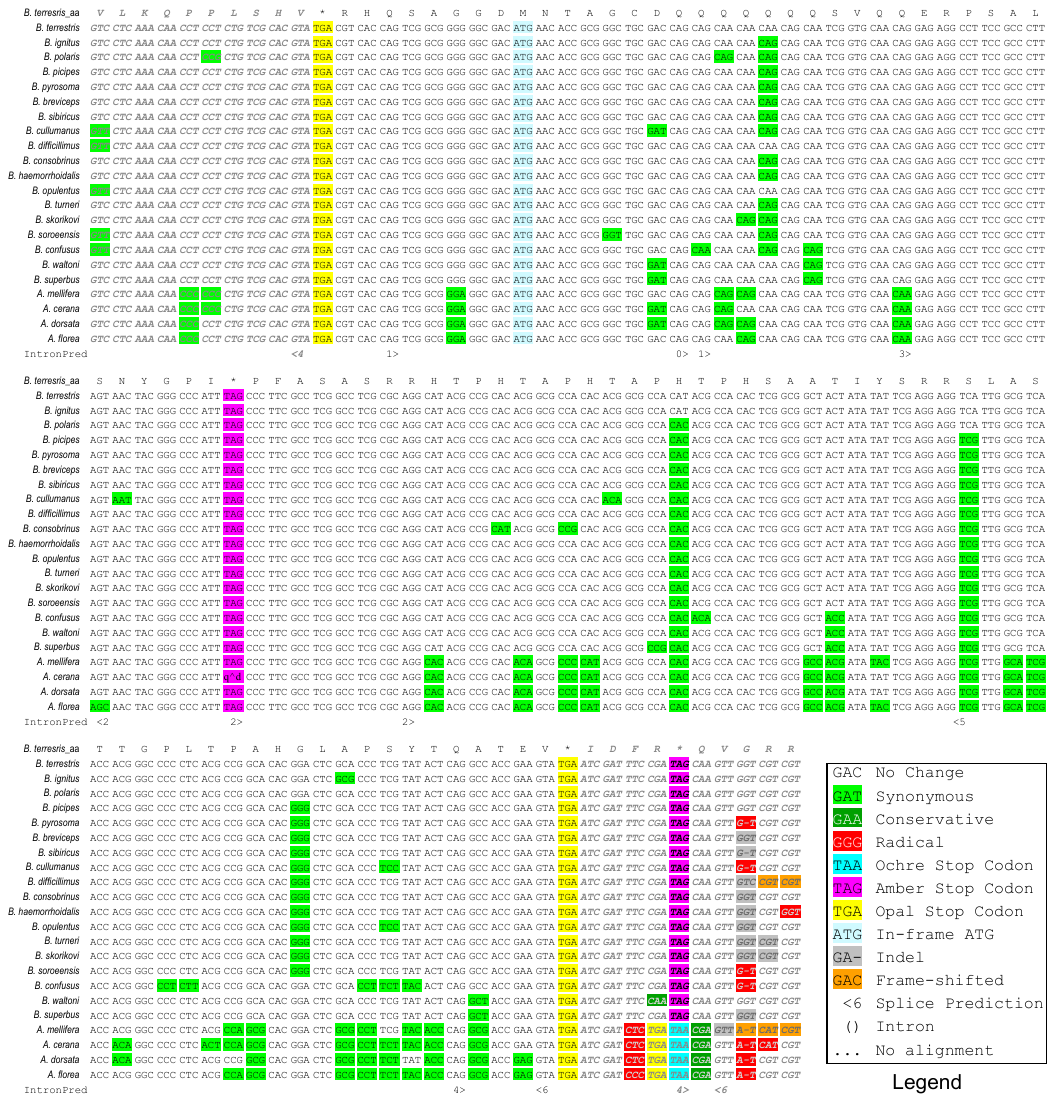

### Figure S13: Tree with nodes labeled for Malin analysis.

Species names in the tree are in short and their corresponding full names are as follow: Bpici (*B. picipes*), Bigni (*B. ignitus*), Bpyro (*B. pyrosoma*), Bturn (*B. turneri*), Bsupe (*B. superbus*), Bsoro (*B. soroeensis*), Bcull (*B. cullumanus*), Bpola (*B. polaris*), Bhaem (*B. haemorrhoidalis*), Bconf (*B. confusus*), Bsibi (*B. sibiricus*), Bcons (*B. consobrinus*), Bopul (*B. opulentus*), Bskor (*B. skorikovi*), Bbrev (*B. breviceps*), Bdiff (*B. difficillimus*), Bwalt (*B. waltoni*), Bterr (*B. terrestris*), Bimpa (*B. impatiens*) , Amell (*Apis mellifera*).

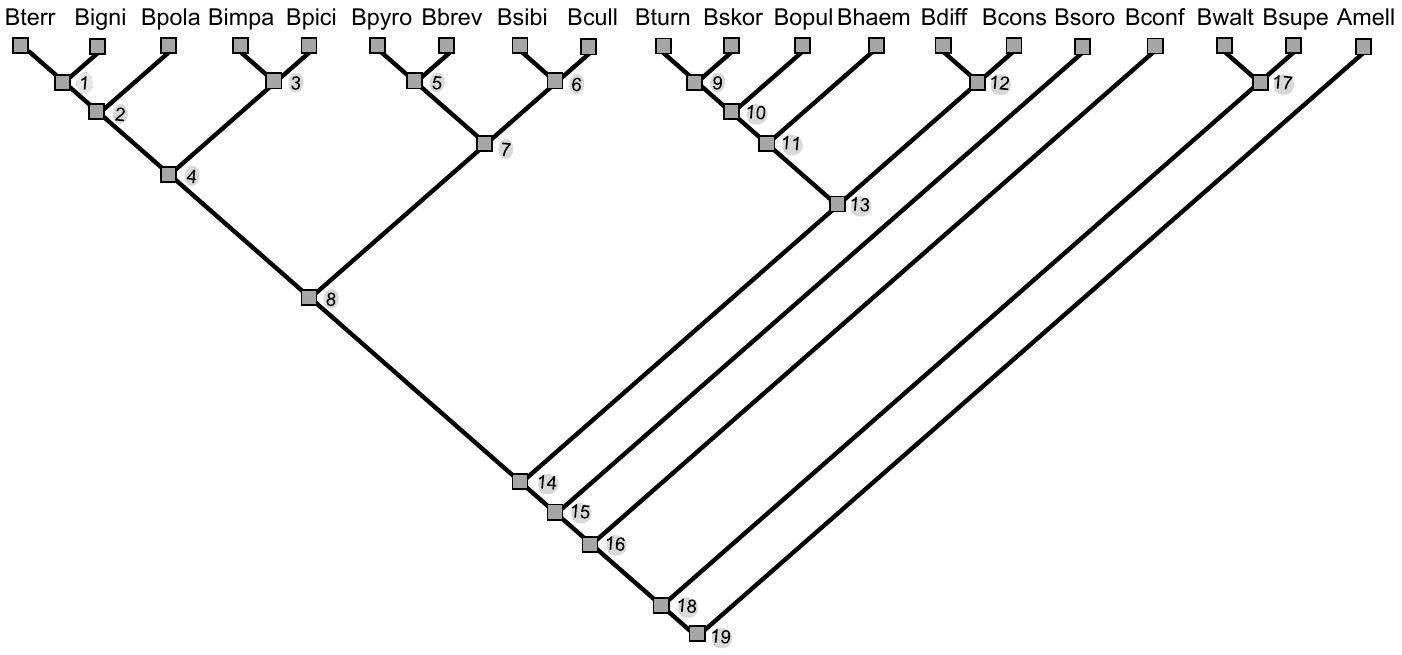

### Figure S14: Functional annotation bias towards conserved genes.

Histograms show value distributions for all orthologous groups and for orthologous groups with genes that could be assigned by (**A**). Biological process GO terms (in green), (**B**). Molecular function GO terms (in green), and (**C**). InterPro domains (in green).

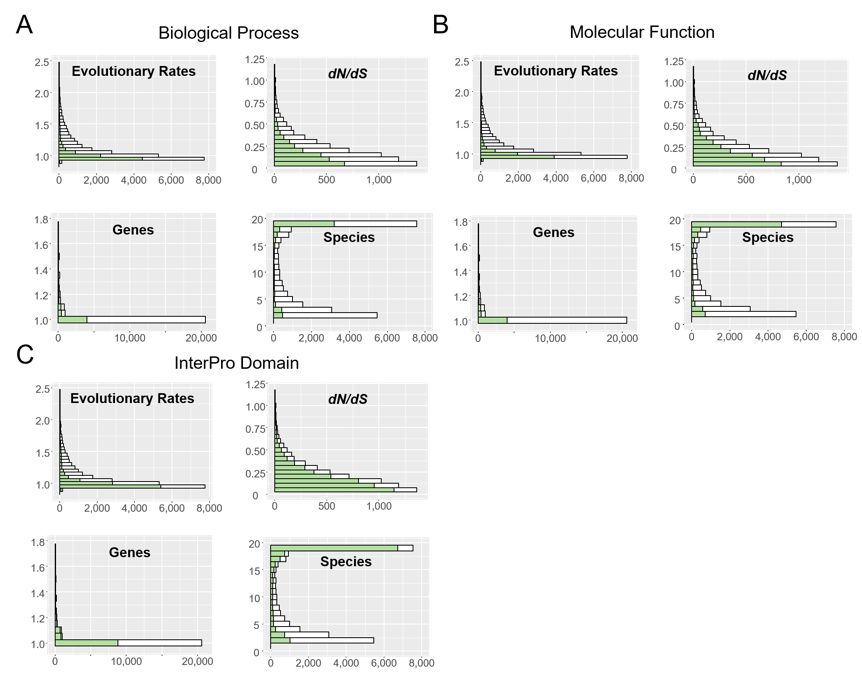

### Figure S15: Molecular evolution of protein-coding genes in term of evolutionary rate (amino acid sequence divergence) and *dN/dS* ratio among gene functional classes categories by (A) Molecular functions GO terms and (B) InterPro domains.

Categories are sorted by evolutionary rate from the most conservative (left) to the most dynamic (right) and colored from the highest values (red) to the median value (blue) to the lowest values (orange). Notched boxes show medians of orthologous group values with the limits of the upper and lower quartiles, and box widths are proportional to the number of orthologous groups in each category.

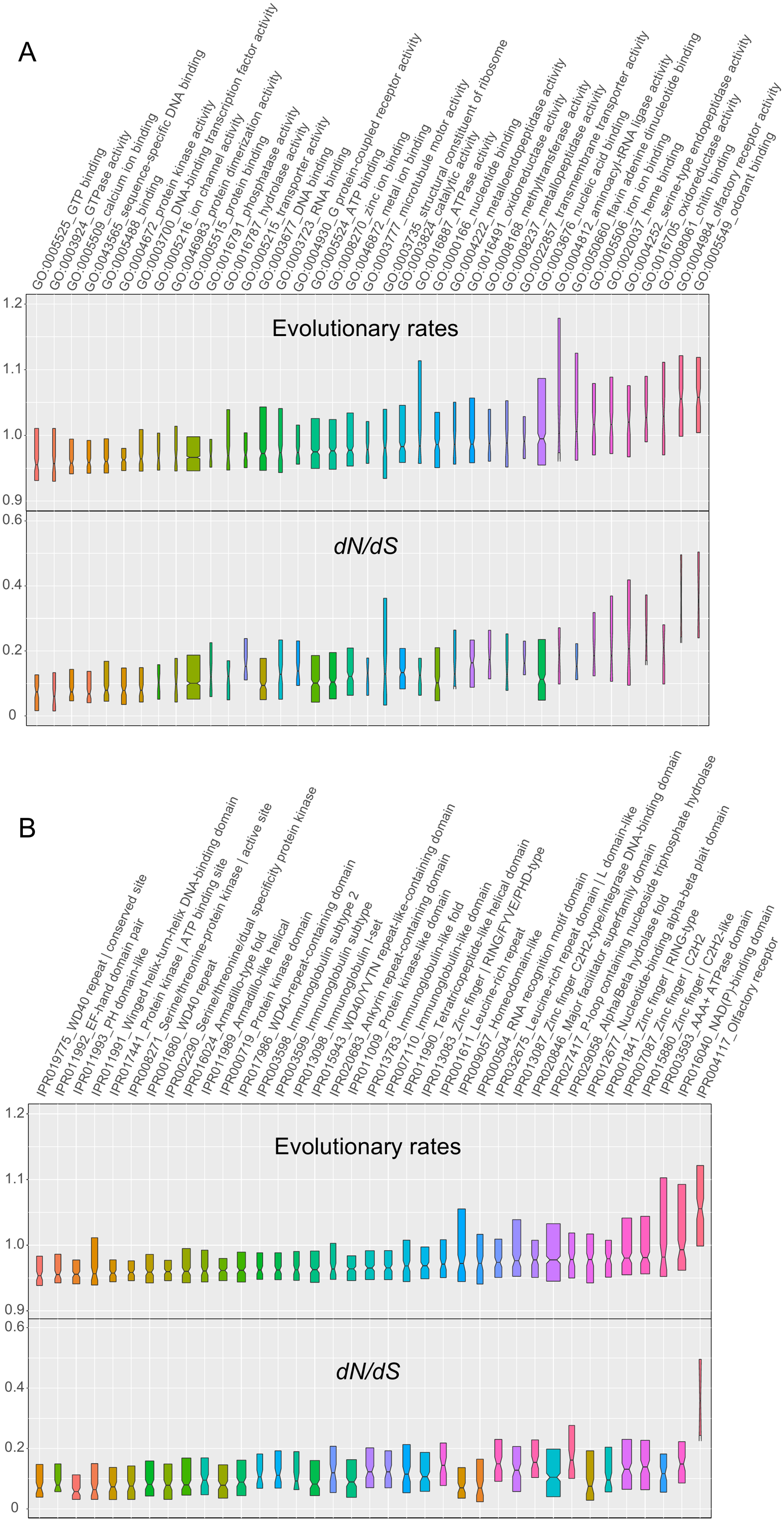

### Figure S16: Evolutionary rate and *dN/dS* ratio distributions.

(**A**). Distribution of orthologous group evolutionary rates highlighting those less than the 20th percentile or greater than the 80th percentile. (**B**). Distribution of orthologous group *dN/dS* ratios highlighting those less than the 20th percentile or greater than the 80th percentile.

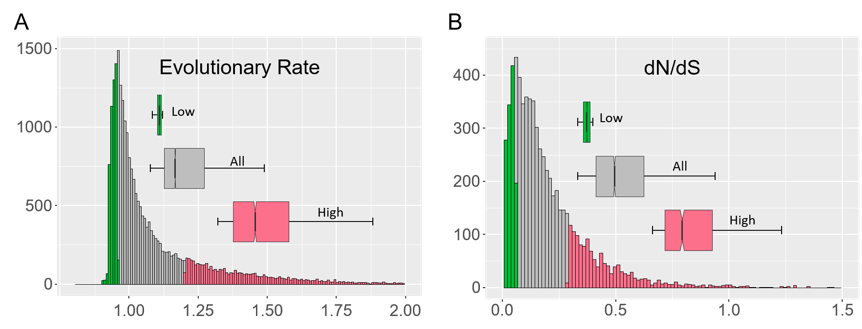

### Figure S17: Correlation coefficient between RSCU and ENC, estimated as exponential of the sum of Shannon entropy of codon usage within each codon family across the 19 species in rows and the 64 codons in columns.

Black lines separate codon family. Blue values indicate negative correlation, meaning preferred codon, given that high frequency of a codon correlates with decrease of entropy in codon usage across the genes and codon family.

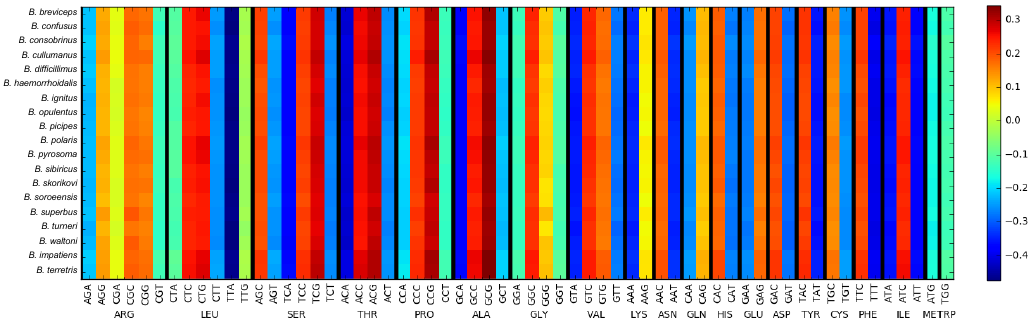

### Figure S18: Correlation between gene AT content and the frequency of optimal codons.

Optimal codons are defined as the most negatively correlated codon within each family (Data from Figure S17). The scale bar on the side indicates number of points in each hexagonal bin.

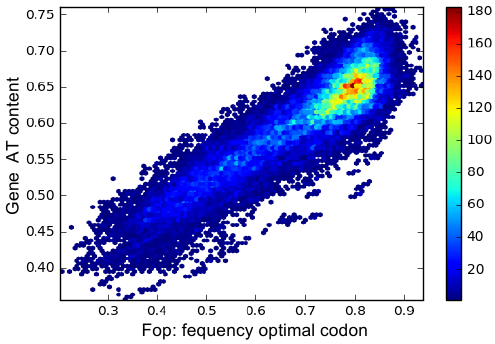

### Figure S19: Relationship between codon AT content and correlation shown in Figure S17.

Numbers of overlapping points within each hexagonal bin are defined in the color bar.

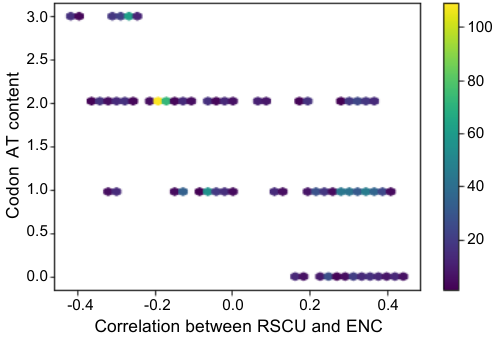

### Figure S20: Evolutionary histories of chemosensory genes in bumblebees.

OR represents odorant receptor; GR represents gustatory receptors; IR represents ionotropic receptors. The results of CAFE and Notung are highlighted in blue and red, respectively. Number of intact chemosensory genes are shown beside species name. Numbers on each node are the estimated ancestral gene numbers. Numbers on branches indicate gene gain and loss events estimated by Notung.

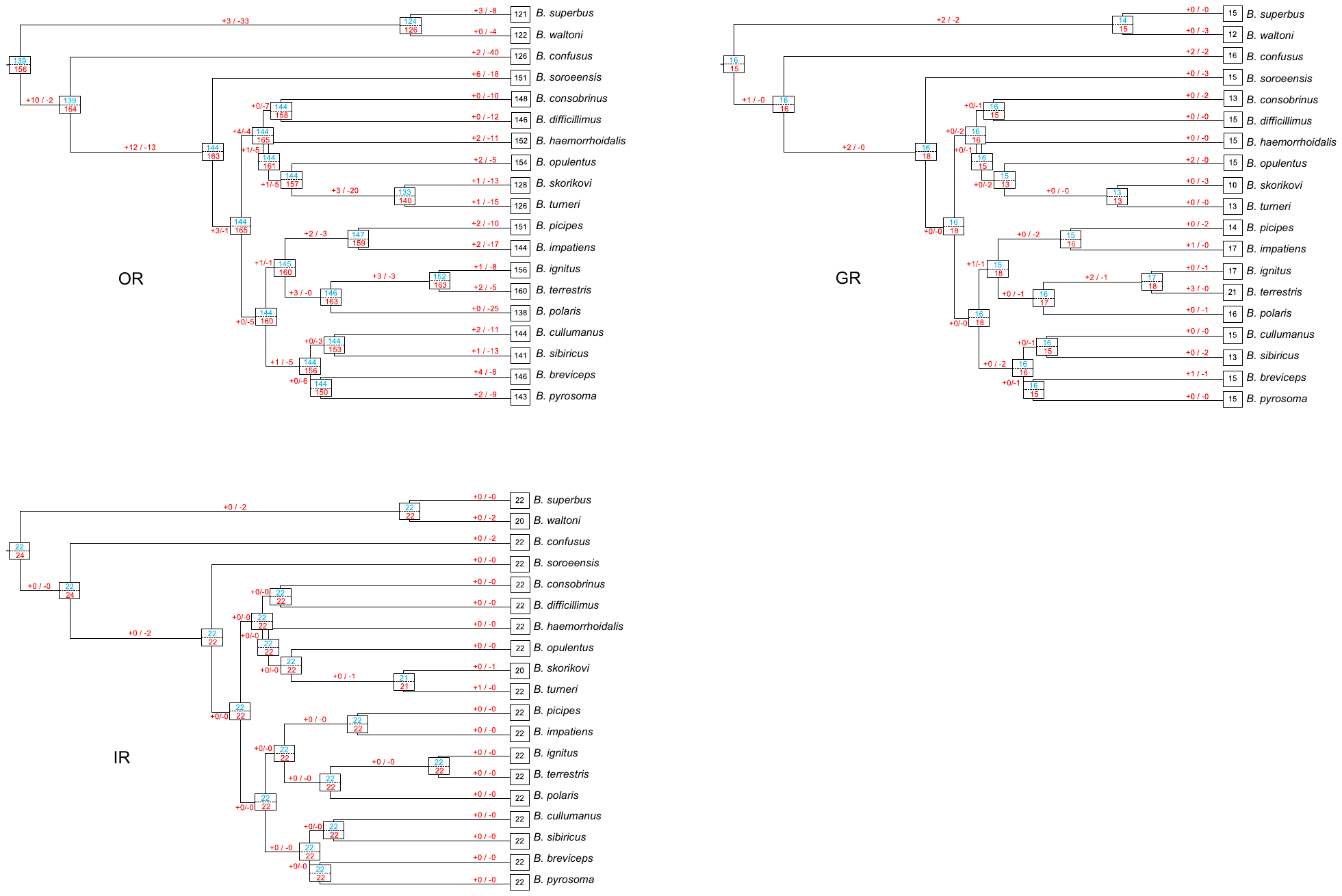

### Figure S21: Sex-determination genes fem and fem1.

**Example of amino acid motif distribution for *fem*, *fem1* (and *csd*) among *Bombus* and *Apis*, with highlighting lineage specific (*fem1*) motifs and taking structural variation into account.**

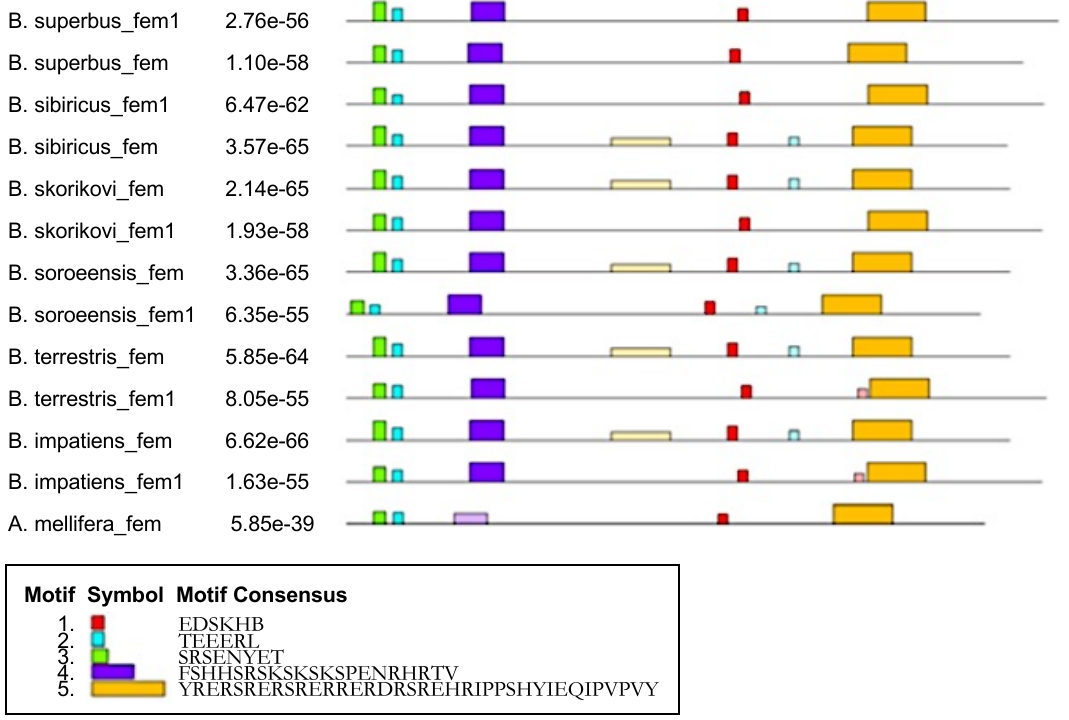

### Figure S22: Sex-determination gene tra2.

**Tra2 RNA recognition domain with RNA binding sites marked (in grey conserved; in red changed in *Apis* compared to *Bombus*).**

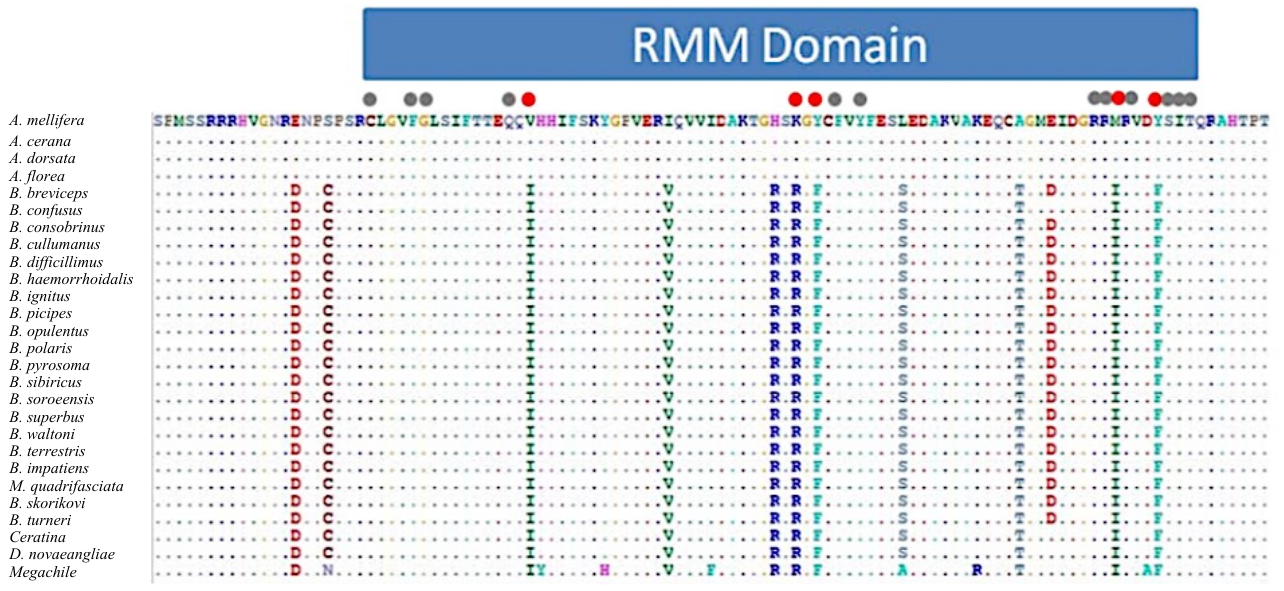

### Figure S23: Genome assembly sizes versus contiguity.

**Pearson correlation analysis between genome assembly sizes and genome assembly contiguity (scaffold N50).**

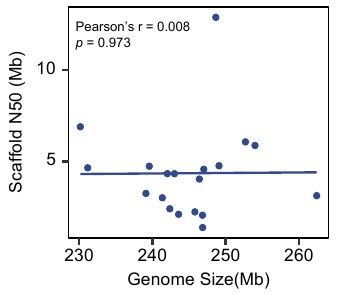
